# Saturation Genome Editing Resolves the Functional Spectrum of Pathogenic *VHL* Alleles

**DOI:** 10.1101/2023.06.10.542698

**Authors:** Megan Buckley, Christina M. Kajba, Nicole Forrester, Chloé Terwagne, Chelsea Sawyer, Scott T.C. Shepherd, Joachim De Jonghe, Phoebe Dace, Samra Turajlic, Gregory M. Findlay

## Abstract

To maximize the impact of precision medicine approaches, it is critical to accurately identify genetic variants in cancer-associated genes with functional consequences. Yet, our knowledge of rare variants conferring clinically relevant phenotypes and the mechanisms through which they act remains highly limited. A tumor suppressor gene exemplifying the challenge of variant interpretation is *VHL*. *VHL* encodes an E3 ubiquitin ligase that regulates the cellular response to hypoxia. Germline pathogenic variants in *VHL* predispose patients to tumors including clear cell renal cell carcinoma (ccRCC) and pheochromocytoma, and somatic *VHL* mutations are frequently observed in sporadic renal cancer. Here, we optimize and apply Saturation Genome Editing (SGE) to assay nearly all possible single nucleotide variants (SNVs) across *VHL*’s coding sequence. To delineate mechanisms, we quantify mRNA dosage effects over time and compare effects in isogenic cell lines. Function scores for 2,268 *VHL* SNVs identify a core set of pathogenic alleles driving ccRCC with perfect accuracy, inform differential risk across tumor types, and reveal novel mechanisms by which variants impact function. These results have immediate utility for classifying *VHL* variants encountered in both germline testing and tumor profiling and illustrate how precise functional measurements can resolve pleiotropic and dosage-dependent genotype-phenotype relationships across complete genes.

## INTRODUCTION

Delineating rare genetic variants underlying disease phenotypes remains a major challenge in human genetics. Genome-wide association studies and germline DNA sequencing have advanced our knowledge of genes and pathways implicated in cancer predisposition. Concurrently, genomic profiling of tumors has revealed mutations that frequently arise somatically during cancer evolution, indicating their functional importance. Yet, for the majority of cancer-associated genes, more variants of uncertain significance (VUS) have been reported than variants whose phenotypic effects are known^1–4^. In the context of both germline testing and tumor profiling, VUS represent a missed opportunity to improve patient care through precision medicine approaches.

Most variants are difficult to interpret with high confidence because they are too rare to enable statistically robust genotype-phenotype associations. Computational models of variant effect have improved due to greater availability of training data and the use of machine learning^5–10^. However, such models are not accurate enough to dictate clinical decisions without additional evidence^11^. Mechanistic knowledge of variants in tumor suppressor genes can lead directly to improved patient care by informing which individuals will benefit from preventative measures and by guiding selection of targeted anticancer therapies^12–14^. There is, however, a scarcity of functional data available for linking variants to phenotypes^15^.

The von Hippel-Lindau tumor suppressor (VHL) is a 213-amino acid (AA) protein encoded on chromosome 3p which functions as an E3 ubiquitin ligase in complex with Cullin-2 (CUL2), Elongins C and B (ELOC, ELOB), and ring-box 1 (RBX1)^16^. In normoxic conditions, VHL ubiquitinates the ɑ-subunit of hypoxia-inducible factor (HIF), targeting HIF for proteasomal degradation. In hypoxic conditions, HIF is protected from VHL-mediated degradation and signals to promote glycolysis and angiogenesis. Loss of VHL function due to mutation can lead to constitutive HIF activity and, in certain tissues, tumor development^17^.

Somatic *VHL* mutations are frequently observed in renal cell carcinomas (RCCs), most commonly clear cell RCC (ccRCC). During ccRCC evolution, chromosome 3p deletion typically precedes a loss-of-function (LoF) mutation to the remaining *VHL* allele, resulting in increased HIF activity^18^. *VHL* mutations have been observed in other types of RCC, as well as extrarenal cancers, but their functional significance is less certain. Currently, 185 *VHL* variants are reported to be VUS in the cBioPortal database^1, 2^, including 39% of observed missense variants.

Pathogenic germline variants in *VHL* predispose patients to different neoplasias in an autosomal dominant manner, a rare condition known as VHL disease^19^. Affected patients have varying susceptibility to different tumors including ccRCC, pheochromocytoma, and hemangioblastoma. The risk and frequency of each tumor depend largely upon the specific germline mutation. Classically, type 1 VHL disease variants lead to complete loss of VHL function, and include whole-gene deletions, nonsense variants, frameshifting insertions and deletions (indels), as well as certain splice variants. Type 1 variants predispose patients to ccRCC, as well as hemangioblastomas and other neoplasms in a HIF-dependent manner. Type 2 variants, in contrast, are associated with a high risk of pheochromocytoma and are most often missense variants. Attempts have been made to subclassify type 2 VHL disease further; type 2C disease is marked by pheochromocytomas only, type 2A disease includes hemangioblastomas and other benign tumors, and type 2B disease further includes ccRCC^20^.

These broad, clinical classifications of VHL disease have helped explain patterns of tumors present in families. However, a complete molecular accounting of how different mutations confer distinct pathologies has remained elusive. In a curated database of *VHL* mutations^21^, many variants have been reported to cause both type 1 and type 2 disease. Other *VHL* variants have been implicated in recessive diseases, such as congenital polycythemia or germline VHL deficiency^22, 23^. Although protein-truncating variants typically cause type 1 disease, in rare instances, patients with nonsense variants have presented with type 2 disease marked by early-onset pheochromocytomas^24^. Such discrepancies highlight the limitations of observational approaches and the challenge of developing individualized surveillance and therapy plans for patients without mechanistic knowledge of variants’ effects.

In addition to variants known to cause VHL disease, there are over 800 VUS in *VHL* reported in ClinVar^3^. It is unknown what fraction of these variants cause disease, and likewise how many variants yet to be reported may prove pathogenic. The fact that many variants have been observed repeatedly in unrelated families and across independent cancers suggests genetic evidence may be converging on a near-complete set of pathogenic alleles^21^. Yet, substantially more *VHL* variants are predicted by computational models to be deleterious than have been linked

to disease^25^. It is unclear whether such variants have yet to be encountered due to their rarity, whether they are incompatible with life, or whether they simply do not cause disease.

For patients harboring *VHL* variants whose phenotypic effects are unknown, well-calibrated functional data may prove useful in aiding diagnosis and management. Furthermore, the recently demonstrated efficacy of HIF2A inhibitors for preventing ccRCC progression^14^ suggests quantifying *VHL* variants’ precise effects on HIF regulation may prove valuable for guiding therapeutic selection. More broadly, *VHL* serves as a powerful gene model to assess the extent to which functional data can recapitulate genotype-phenotype relationships in humans, owing to the fact it is nearly always mutated during ccRCC development^2^ and the extensive knowledge regarding phenotypic effects of many germline variants^21^.

Here, we systematically measure the functional consequences of *VHL* variants across the complete gene by employing Saturation Genome Editing (SGE). In total, we score 2,268 SNVs for HIF-dependent effects on cellular fitness, defining LoF variants underlying ccRCC development with 100% accuracy. Our assay captures clinically meaningful differences in the degree of functional impairment among pathogenic alleles and delineates novel mechanisms explaining genotype-phenotype associations in humans, suggesting a role for improving diagnostic and therapeutic precision.

## RESULTS

### A highly optimized SGE assay to precisely measure effects of *VHL* variants

Towards developing a high-throughput assay to study variants in *VHL*, we analyzed data from genome-wide knockout screens to assess the effect of *VHL* loss across cell lines. With the exception of kidney-derived lines, CRISPR-induced knockout of *VHL* almost uniformly reduces cell fitness (**Fig. 1a**)^26, 27^. We reasoned growth defects may, therefore, be a scalable means of assaying *VHL* variants for LoF effects.

**Fig. 1:**
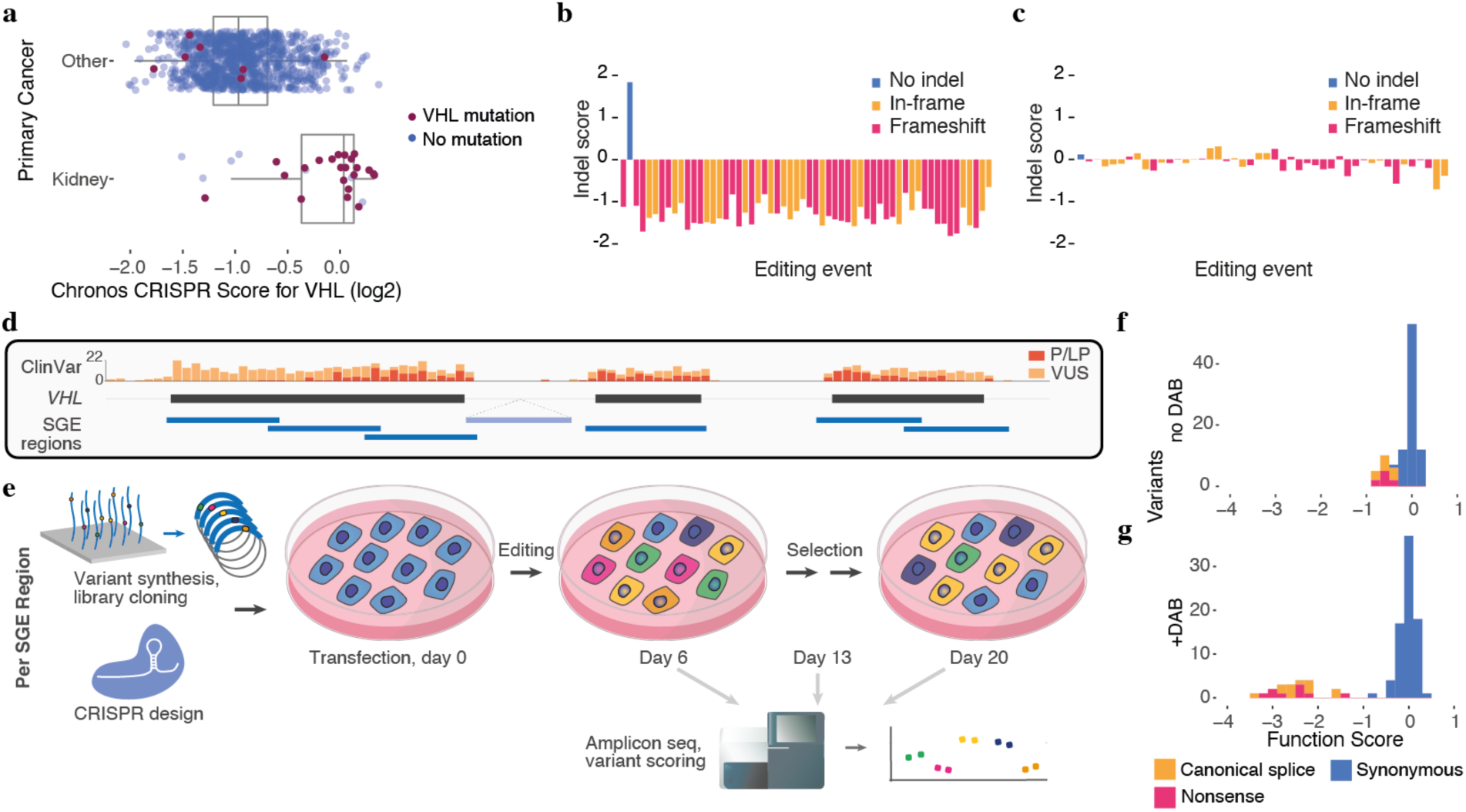
A highly optimized SGE protocol to assay *VHL* variants. **a**, CRISPR knock-out screening data from the Cancer Dependency Map (DepMap)^26^ reveal *VHL* loss widely leads to reduced growth in cell lines lacking *VHL* mutations (boxplot: center line, median; box limits, upper and lower quartiles; whiskers, 1.5x interquartile range; all points shown). **b,c**, CRISPR-induced editing of *VHL* was performed in HAP1 cells (day 0) and outcomes were quantified by next-generation sequencing (NGS). Indel scores, calculated for each editing outcome as the log_2_-ratio of abundance on day 13 to day 6, are shown, ranked by day 6 frequency. Frameshifting indels are strongly selected against in parental HAP1 (**b**) but not HIF1A-knockout cells (**c**). **d**, The strategy to perform SGE across the complete coding sequence of *VHL* is shown, with ClinVar variant counts for “pathogenic” and ”likely pathogenic” variants (red) and VUS (orange) displayed from gnomAD^36^. SGE regions were designed to tile exons 1-3, as well as a region of intron 1. **e**, For each SGE region, a library of oligos containing all possible SNVs was synthesized and cloned into a vector with homology arms to facilitate genomic integration via CRISPR-induced homology-directed DNA repair (HDR). Variants present in cells were quantified over time via amplicon sequencing and function scores were calculated to reflect variants’ effects on fitness. **f,g**, Function scores for synonymous, nonsense, and canonical splice site SNVs are shown for a single SGE region (exon 2) assayed in normal media (**f**) or media supplemented with 2.5 µM DAB (**g**).

To investigate *VHL*’s essentiality in the haploid human line HAP1, indels in exon 2 were generated with CRISPR and sequenced at multiple timepoints following editing. Robust depletion of frameshifting indels compared to indel-free editing outcomes confirmed the essentiality of *VHL* for normal HAP1 proliferation (**Fig. 1b**). The growth defect upon targeting *VHL* with CRISPR was reduced by simultaneous targeting of *HIF1A* (**Supplementary Fig. 1**), and selection against indels was effectively eliminated by prior knockout of *HIF1A* (**Fig. 1c**). Collectively, these data show *VHL* loss confers a HIF-dependent growth defect in HAP1 cells.

SGE is a method by which all possible SNVs in a genomic region of up to 150 bp are assayed in multiplex using CRISPR/Cas9 editing^28^. When SGE is performed in HAP1, a single variant is engineered per haploid cell. This allows variants’ effects on growth to be accurately determined by NGS. Seven SGE libraries were made to cover the full coding sequence of *VHL*, as well as exon-proximal regions of introns and a region deep within intron 1 (**Fig. 1d**). Each library consisted of all possible SNVs in a region cloned into vectors with homology arms to facilitate genomic integration (**Fig. 1e**). Sequencing of initial libraries revealed two regions of exon 1 to have skewed variant distributions at sites of repetitive, GC-rich sequence (**Supplementary Fig. 2a-e**). Additional synonymous mutations were therefore engineered in these regions, resulting in improved library uniformity (**Supplementary Fig. 2f-l**).

With the aim of measuring more subtle effects on growth, SGE experiments were performed using a protocol modified from published work^29^ featuring optimized transfection efficiency and a longer time course (**Fig. 1e**, **Methods**). For each region, replicate transfections were performed on day 0, and approximately 10 million cells were harvested on days 6, 13, and 20. Amplicon sequencing of genomic DNA (gDNA) from each timepoint was used to calculate a “function score” for each SNV reflective of cellular fitness.

For four exonic regions initially assayed, only modest growth defects were observed for expected LoF variants (**Fig. 1f**, **Supplementary Fig. 3**). HAP1 cells can revert to diploidy with prolonged culture^30^, a phenomenon that could weaken recessive effects measured in multiplex. Recently, 10-deacetyl-baccatin-III (DAB) was identified via small molecule screening to select for haploid cells^31^. Therefore, we next performed SGE for all *VHL* regions in media containing 2.5 µM DAB. This led to a marked improvement in dynamic range (**Fig. 1g**, **Supplementary Fig. 3**). In exon 2, for example, the median function score of nonsense and canonical splice site SNVs dropped 4-fold, from -0.62 to -2.49.

We determined SNVs with significantly reduced function scores (i.e. “LoF SNVs”) by defining a null distribution for each SGE region and applying a false discovery rate (FDR) of 0.01. Across SGE regions assayed with and without DAB, there were 39.3% more LoF SNVs identified in DAB-treated cells. After stringent quality filtering, final function scores for *n* = 2,268 SNVs were obtained using data from DAB-treated cells, comprising over 85% of SNVs included in library design (**Supplementary Table 1**).

### Mapping loss-of-function variants

The majority of variants designed but not scored map to the GC-rich 5’ region of exon 1, where the rate of CRISPR editing was lowest (**Supplementary Fig. 4a**). In contrast to other coding regions, the 5’ region of exon 1 did not harbor significantly depleted SNVs, including nonsense variants (**Supplementary Fig. 4b,c**). These data are consistent with a previously characterized alternative translation initiation site at p.M54 producing a fully functional VHL isoform^32^, and corroborate the lack of reported pathogenic variants proximal to p.M54 in ClinVar (**Fig. 1d**).

Moving forward, we restricted analysis to *n* = 2,200 SNVs scored with high reproducibility by excluding SNVs assayed in the 5’-most SGE region of exon 1 (**Fig. 2a,b**). Among *n* = 115 remaining SNVs upstream of p.M54, none were classified as LoF (median score = 0.03). Likewise, no SNVs assayed in the 3’-untranslated region (UTR) or the region deep within intron 1 scored as LoF (**Fig. 2b,c**). In contrast, between p.M54 and p.R200, all but one nonsense variant scored as LoF (*n* = 43, median score = -2.4), as did all canonical splice site SNVs (*n* = 24, median score = -2.3) (**Fig. 2b,c**). Most missense variants scored neutrally, though 22.4% were scored as LoF.

**Fig. 2:**
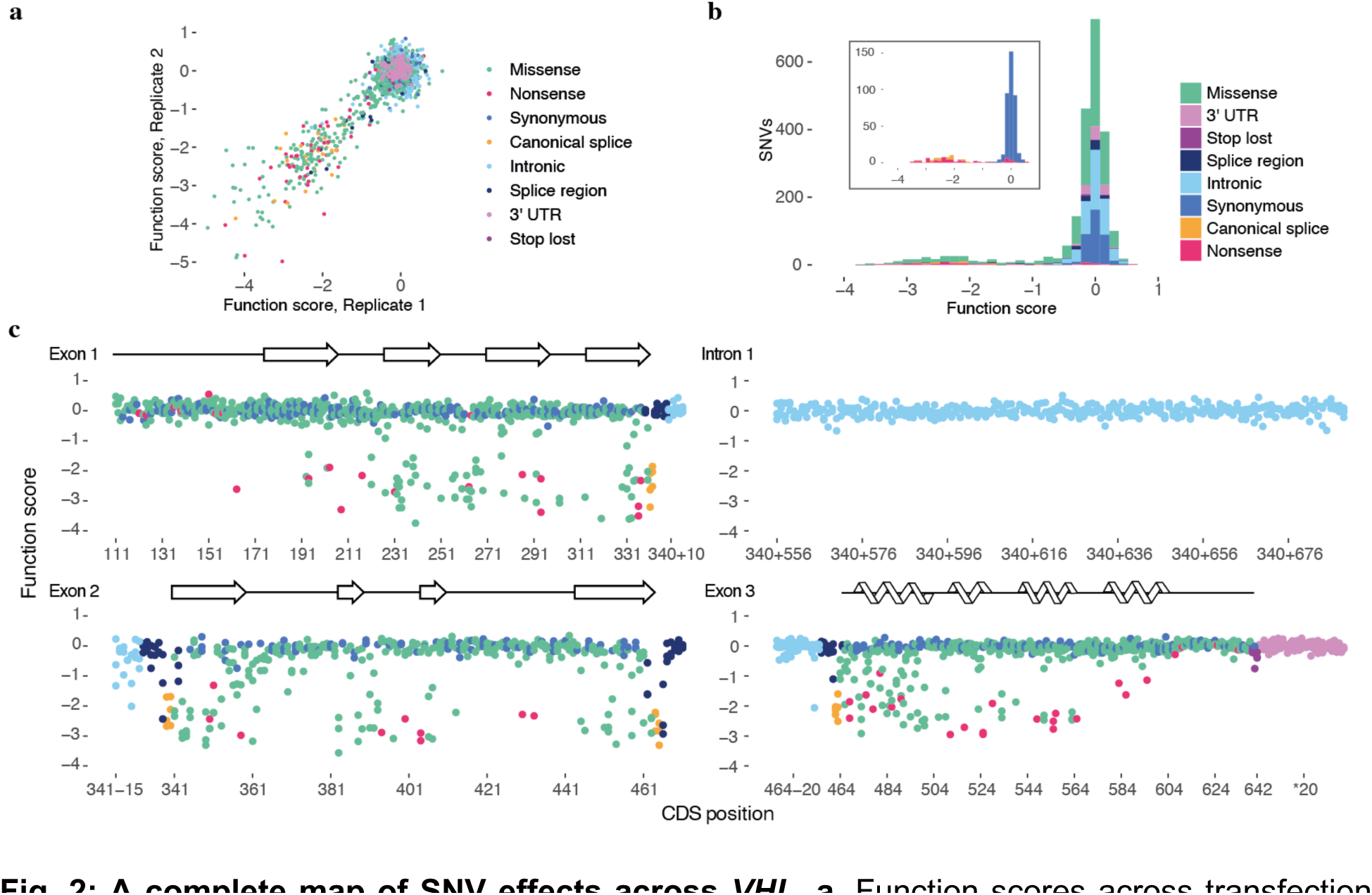
A complete map of SNV effects across *VHL*. **a**, Function scores across transfection replicates are plotted for *n* = 2,200 SNVs (Pearson’s *R* = 0.90). **b**, Replicate scores were averaged and normalized to obtain a final function score for each variant. The inset shows only synonymous, nonsense, and canonical splice site SNVs. **c**, Function scores are plotted by genomic position for each coding and intronic region assayed, with β-sheets and ɑ-helices of VHL’s secondary structure^33^ shown above.

Common mechanisms by which variants can lead to LoF include reducing expression of mRNA (for instance, by disrupting splicing) and altering protein function (for instance, by causing instability or loss of enzymatic activity). To better resolve mechanisms of functional impairment, we derived *n* = 1,626 “RNA scores” for coding variants by performing targeted RNA-sequencing of day 6 and day 20 samples (**Supplementary Fig. 5, Supplementary Table 1**). RNA scores reflect SNVs’ effects on full-length *VHL* mRNA levels.

Comparison of RNA scores to function scores reveals only large reductions in mRNA confidently predict LoF (**Fig. 3a**). Indeed, 94% of SNVs with RNA scores below -3 scored as LoF (*n* = 17, function scores from -2.9 to -0.38), suggestive of a minimum mRNA dosage required for normal cell growth. Many SNVs with low RNA scores occur near exon-intron junctions, where variants are more likely to disrupt splicing (**Supplementary Fig. 5**). While RNA scores across timepoints were highly correlated, variants strongly depleted in mRNA on day 6 tended to be less depleted in mRNA on day 20, consistent with selective effects on transcript levels (**Supplementary Fig. 6**).

**Fig. 3:**
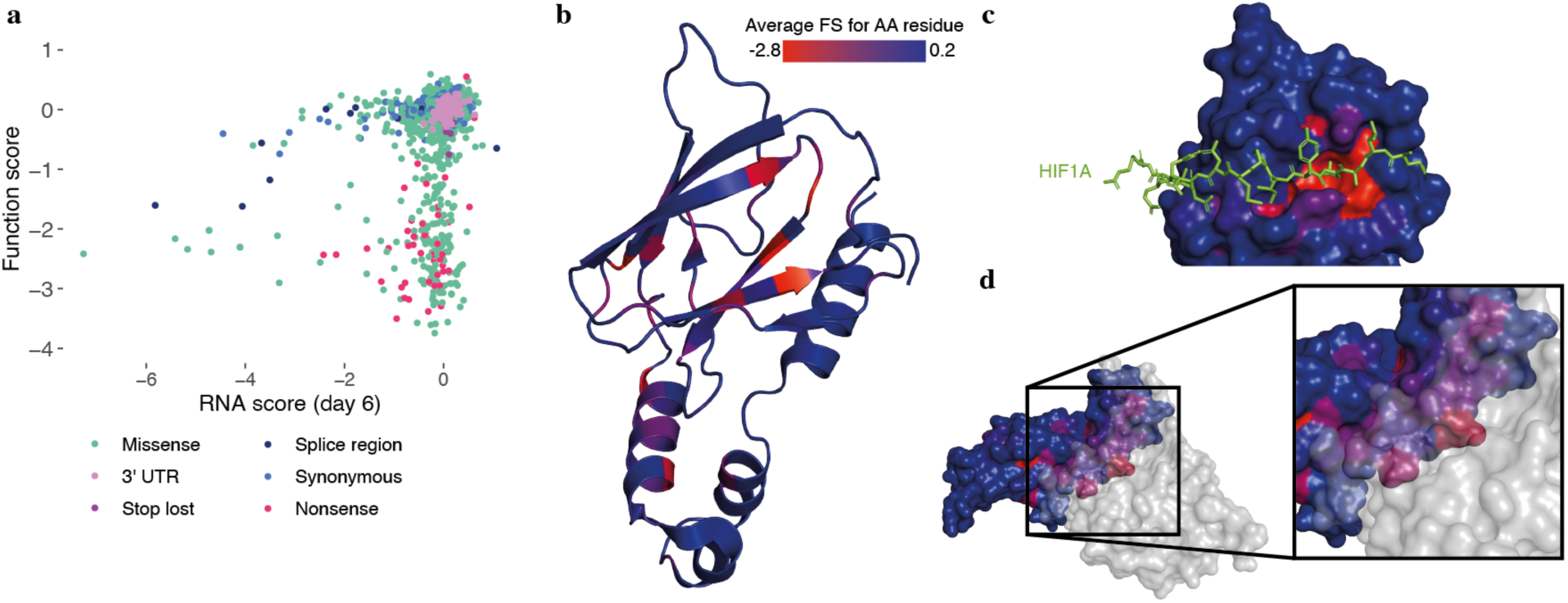
Function scores capture fitness effects mediated at both the transcript and protein level. **a**, Targeted RNA-seq of *VHL* mRNA from edited cells was performed to calculate RNA scores for *n* = 1,626 exonic SNVs. The relationship between RNA scores and function scores reveals most LoF variants are expressed at normal levels in mRNA, and only low RNA scores reliably predict LoF at the cellular level. **b-d**, The average score of missense variants at each amino acid position is shown on the VHL structure (PDB: 1LM8)^47^. Positions intolerant to variation include residues within β-sheets (**b**), residues forming the HIF1A recognition site (**c**), and contacts at the ELOC interface (**d**).

The majority of LoF SNVs, however, did not have low RNA scores. This suggests their effects are mediated at the protein level. Of note, nonsense variants as a class were only minimally depleted in mRNA (median RNA score = -0.19, **Supplementary Fig. 5**), indicating minimal nonsense-mediated decay (NMD). Overlaying function scores to the VHL structure^33^ reveals missense variants that score lowly tend to occur in β-sheets and ɑ-helices (**Fig. 3b**). Residues highly intolerant to missense variation include those forming the substrate recognition site as well as specific contacts with ELOC (**Fig. 3c,d**). Collectively, these findings confirm most LoF variants exert effects by altering function at the protein level, with only large decreases in *VHL* mRNA sufficient to cause functional impairment in the assay.

### Function scores accurately distinguish *VHL* variants driving human disease

To assess whether function scores predict variants driving human disease we performed several analyses. First, using all *VHL* variant annotations in ClinVar meeting assertion criteria (i.e. awarded at least one star), we observed that function scores distinguish “pathogenic” variants from “benign” and “likely benign” variants with 95.2% sensitivity and 97.9% specificity (**Fig. 4a**, **Supplementary Fig. 7a**). Performance is slightly worse upon inclusion of “likely pathogenic” variants (**Supplementary Fig. 7b,c**). Importantly, this broad analysis includes variants with diverse phenotypes described in ClinVar.

**Fig. 4:**
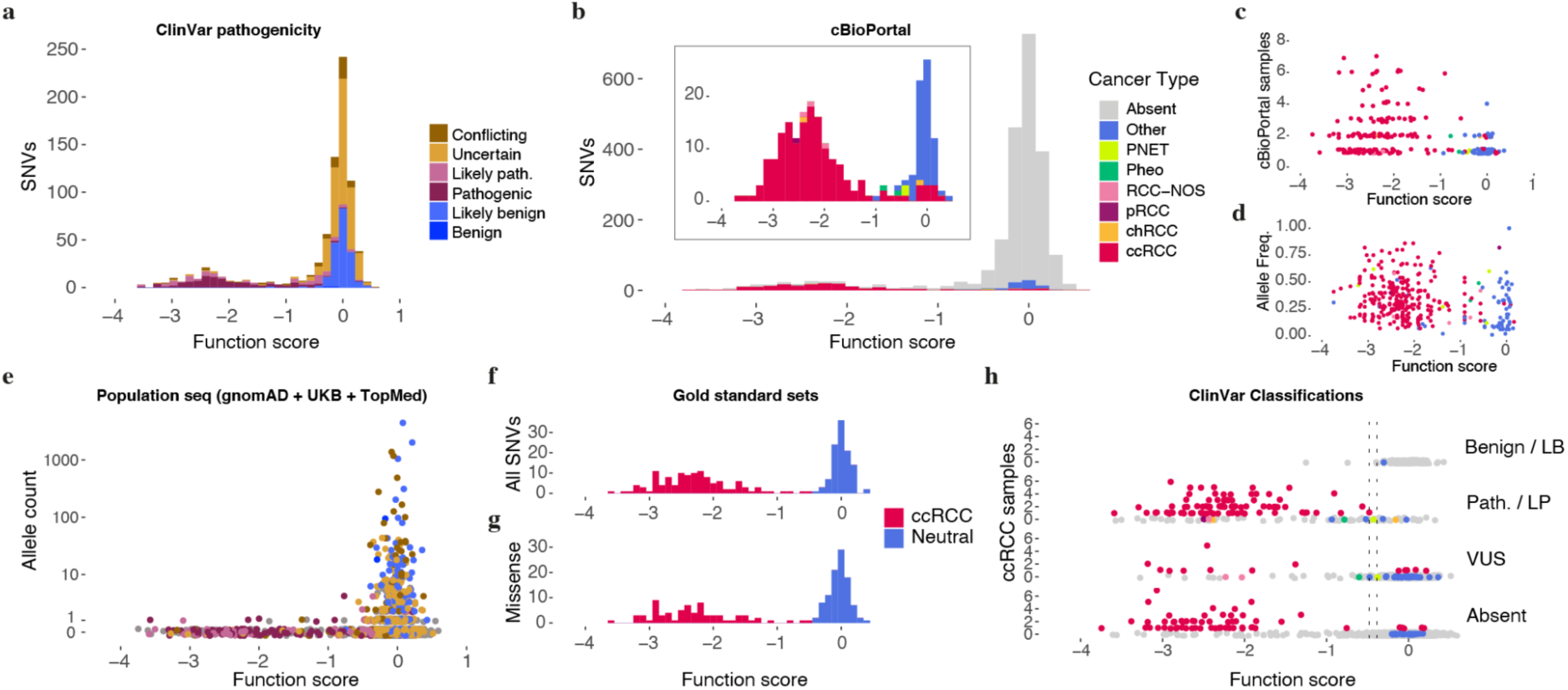
Highly accurate identification of pathogenic *VHL* alleles in humans using SGE. **a,b** The distribution of function scores for *VHL* SNVs reported in ClinVar^3^ (**a**) and observed in cBioPortal^1, 2^ cancer samples (**b**) are shown. **c**,**d**, For SNVs observed in tumors, lower function scores correlate with more independent observations in cBioPortal (**c**) (*n* = 233 SNVs, Spearman’s ⍴ = 0.25, *P* = 9.7 x 10^-5^) and higher allele frequencies within samples (**d**) (*n* = 334 samples, ⍴ = 0.22, *P* = 3.7 x 10^-5^). **e**, A combined population allele count for each SNV assayed was determined by summing independent observations from gnomAD^36^, UK Biobank^35^, and TOPMed^37^. Variants observed repeatedly in population sequencing nearly always scored neutrally. **f,g**, Gold-standard sets of variants were defined using orthologous data from ClinVar, cBioPortal, and population sequencing. In **f**, all variants with at least 2 cBioPortal ccRCC observations or 1 ccRCC observation and a ClinVar “pathogenic” or “likely pathogenic” annotation (*n* = 120 ccRCC-associated SNVs) are plotted with variants deemed “benign” or “likely benign” in ClinVar and seen at least once in population sequencing (*n* = 108 neutral SNVs). **g**, Due to the lack of missense variants classified as “benign” or “likely benign” in ClinVar, we defined a neutral set to include those present in at least 2 population controls and not deemed “pathogenic” or “likely pathogenic” in ClinVar (*n* = 99 neutral missense SNVs versus *n* = 73 ccRCC missense SNVs). **h**, Function scores are plotted against cBioPortal ccRCC observations for SNVs reported in ClinVar to be “benign” or “likely benign” (LB), “pathogenic” or “likely pathogenic” (LP), or VUS (here, including SNVs with conflicting interpretations), and for SNVs absent from ClinVar. SNVs are colored by cancer type (as in **b**). Lines correspond to thresholds for pathogenicity (-0.479) and benignity (-0.388) that delineate gold-standard ccRCC variants in **f**.

Next, tumor sequencing data from cBioPortal^1, 2^ was used to examine the functional effects of *VHL* mutations seen across human cancers. Over 93% of SNVs seen in at least one RCC sample of any type scored as LoF (**Fig. 4b**). This was in contrast to *VHL* variants in other tumor types, which typically scored neutrally, with the exception of variants in pheochromocytomas and pancreatic neuroendocrine tumors (PNETs) – extrarenal tumors also linked to germline *VHL* mutations^34^. SNVs observed in more individual patients and at higher allele frequencies had significantly lower function scores, confirming LoF variants defined via SGE are likely driving oncogenesis (**Fig. 4c,d**).

In contrast to most SNVs observed in tumor profiling, nearly all *VHL* SNVs present in population sequencing databases scored neutrally (**Fig. 4e**). For example, among *n* = 119 SNVs seen at least five times in total across the UK Biobank^35^, gnomAD^36^, and TOPMed^37^ databases, no SNV scored below -0.40 (mean = -0.03, s.d. = 0.14). Likewise, the lowest function score for any SNV seen at least twice was -0.77. The narrow distribution of function scores around zero indicates that the vast majority of *VHL* variants seen repeatedly in population sequencing are unlikely to cause VHL disease.

Considering the inherent uncertainty in clinical variant classification and the fact that any mutation may arise in a tumor by chance (i.e. be a “passenger”), we defined a “gold-standard” set of *n* = 120 ccRCC-associated variants supported by multiple lines of evidence. This set includes variants seen in at least two independent ccRCC samples, as well as variants seen in a single ccRCC sample if also deemed “pathogenic” or “likely pathogenic” in ClinVar. SGE function scores perfectly separate these ccRCC-associated SNVs from *n* = 108 SNVs deemed “benign” or “likely benign” in ClinVar and encountered in population sequencing (**Fig. 4f**). Of note, ClinVar lacks “benign” and “likely benign” missense variants in *VHL*. However, function scores also cleanly separate the *n* = 73 missense SNVs in the gold-standard ccRCC set from *n* = 99 missense SNVs not deemed “Pathogenic” or “Likely pathogenic” in ClinVar and seen in population sequencing at least twice (**Fig. 4g**). Collectively, these analyses indicate SNVs scoring below a threshold of -0.478 likely promote ccRCC development, whereas SNVs scoring above -0.388 likely do not.

We next applied these thresholds to assess how function scores may aid clinical classification, both in the context of germline testing and tumor profiling. Of *n* = 430 SNVs reported as VUS or with conflicting interpretations in ClinVar, only 32 (7.4%) scored below -0.478 (**Fig. 4h**). A comparable fraction of SNVs absent from ClinVar scored lowly (132 of *n* = 1,406 SNVs, or 9.4%). Notably, many of these variants have been seen in ccRCC samples, some multiple times. Among SNVs absent from ClinVar that scored below -0.478, those observed in ccRCC tended to score more lowly (median score = -2.51 for ccRCC SNVs vs. median score = -1.21 for SNVs not seen in ccRCC; Wilcoxon rank-sum test *P* = 7.2 x 10^-10^). This indicates that among low-scoring variants, the degree of functional impairment measured by SGE may predict the likelihood of ccRCC development once a mutation arises.

Applying the same thresholds to variants observed in cBioPortal reveals 28 of *n* = 83 SNVs not currently deemed “Oncogenic” likely promote cancer development (**Supplementary Fig. 7d**). Most of these 28 were identified in at least one ccRCC sample. Conversely, of *n* = 150 SNVs currently deemed “Oncogenic,” 11.3% scored neutrally. Such SNVs were often identified in tumors not associated with *VHL* mutations, consistent with these variants not driving disease (**Supplementary Fig. 7d**). In light of these findings, SGE data have clear potential for improving interpretation of variants seen in cancer.

Lastly, we compared function scores for missense variants to outputs from commonly used computational predictors, including CADD^25^, REVEL^5^, EVE^7^, boostDM^8^, and VARITY^9^. Overall, SGE scores performed substantially better at predicting pathogenic missense variants in ClinVar, as well as missense variants in the gold-standard ccRCC set (**Supplementary Fig. 8a,b**). Many missense SNVs scored neutrally by SGE are predicted to be deleterious by the models (**Supplementary Fig. 8c,d**). The absence of such discordantly scored variants from the gold-standard ccRCC set indicates the computational models lack specificity for this phenotype. However, there is generally broad agreement across models in support of variants deemed LoF by SGE.

### A gradient of HIF deregulation implicates novel genetic mechanisms underlying clinical phenotypes

Frameshifting indels in *VHL* were efficiently selected against in HAP1 cells in a HIF1A-dependent manner (**Fig. 1b,c**). To ask whether function scores for SNVs might partially reflect HIF-independent effects, we repeated SGE experiments for SNVs in exons 2 and 3 in HIF1A-knockout cells (*n* = 797 SNVs). All variant effects were effectively eliminated by loss of HIF1A, with there being no correlation between function scores measured across isogenic lines (**Fig. 5a,b**). This finding indicates function scores specifically reflect effects of *VHL* variants on HIF regulation.

**Fig. 5:**
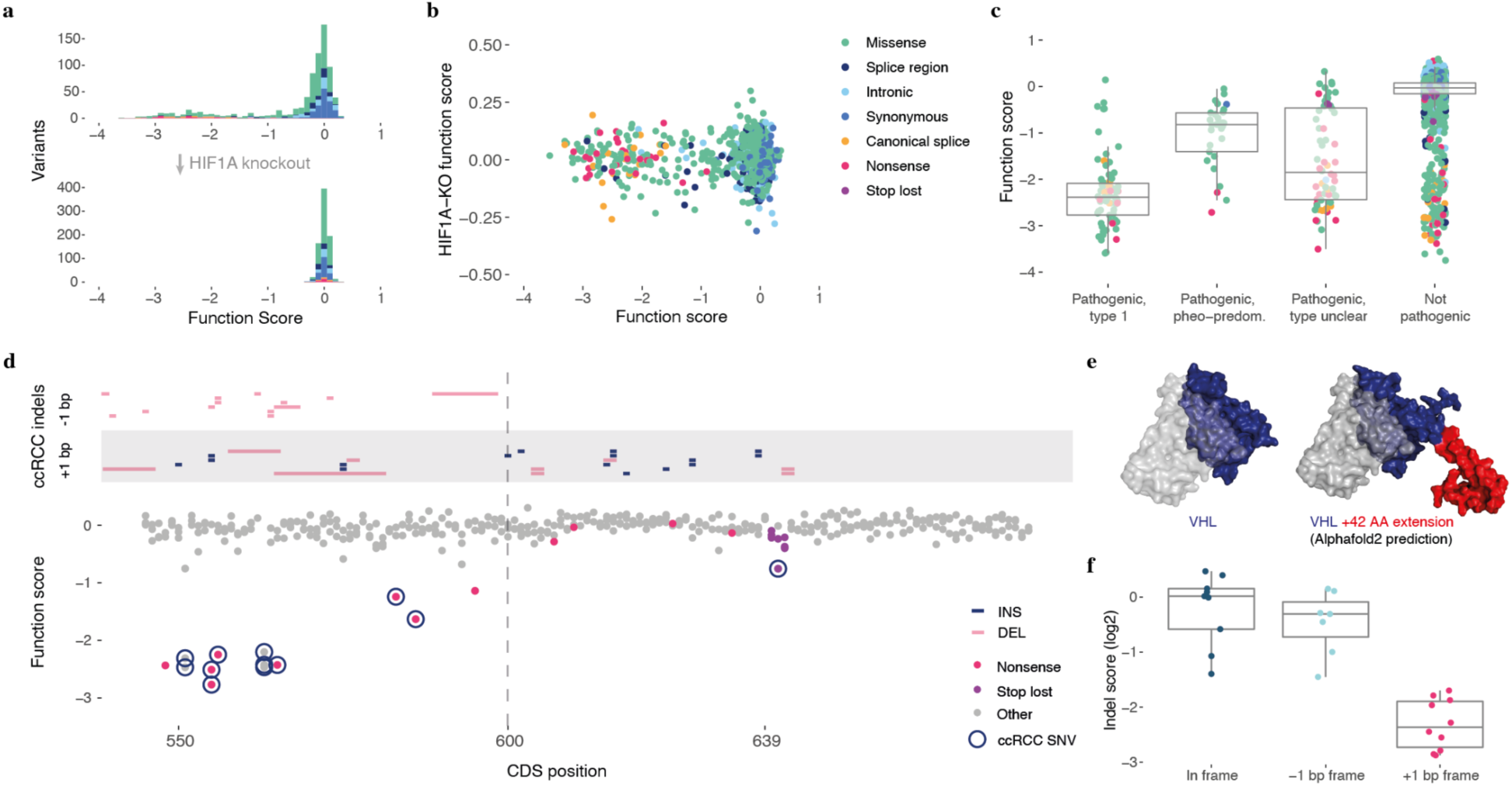
New insights into mechanisms underlying genotype-phenotype associations. **a,** *n* = 797 SNVs in exons 2 and 3 were assayed in HIF1A-KO HAP1 cells. Compared to previous data (top), variants were well-tolerated in HIF1A-KO cells independent of consequence. **b,** Function scores across isogenic lines showed no significant correlation (ρ = -0.06, *P* = 0.08). **c**, “Pathogenic” and “likely pathogenic” SNVs from ClinVar were grouped based on annotations in VHLdb^21^. SNVs associated only with type 1 VHL disease or ccRCC were deemed “type 1” (*n* = 74), whereas SNVs associated only with type 2 disease or predominantly pheochromocytoma were deemed “pheo-predominant” (*n* = 29, excluding SNVs associated with type 2B disease). The remaining pathogenic SNVs lacked unambiguous phenotypic data in VHLdb (*n* = 64, “type unclear”). The boxplot shows function scores for SNVs in each of these categories, as well as for all other SNVs assayed (*n* = 2,033; center line, median; box limits, upper and lower quartiles; whiskers, 1.5x interquartile range; all points shown) **d**, Frameshifting indels present in cBioPortal ccRCC samples are grouped by reading frame and plotted above SGE function scores in the 3’-most region of exon 3. From p.R200, all frameshifting indels seen in ccRCC result in the same reading frame, which extends the protein by 42 AA. **e**, The crystal structure of VHL in complex^47^ is shown next to an AlphaFold prediction^48^ of the full length VHL protein plus the 42-AA C-terminal extension common to indels observed in ccRCC. **f**, Indel scores for CRISPR-induced edits located between p.R200 and p.*214 are plotted, with indels grouped by reading frame. Indels resulting in a +1 bp frameshift scored significantly lower than in-frame indels and -1 bp frameshifting indels (1-way ANOVA: *P* = 1.0 x 10^-7^ and *P* = 2.6 x 10^-6^, respectively; boxplot as above).

Previous studies have shown that certain germline variants conferring high pheochromocytoma risk cause less HIF upregulation^38^. Therefore, we grouped reported pathogenic variants by phenotypic annotations from the VHL Database (VHLdb)^21^. Variants associated only with type 1 disease or ccRCC were deemed “type 1” SNVs, whereas variants associated only with type 2 disease or predominantly pheochromocytomas were deemed “pheo-predominant” SNVs. Critically, the median function score for type 1 SNVs was 2.9-fold lower than for pheo-predominant SNVs (**Fig. 5c**). Pathogenic variants not classifiable in this manner spanned a range of scores, as did LoF SNVs not deemed pathogenic in ClinVar. This result confirms that variants causing high pheochromocytoma risk typically impair HIF regulation to a lesser extent than variants associated only with features of type 1 disease. More broadly, this illustrates how well-calibrated SGE data can inform differential tumor risk across tissues.

In light of the SGE data’s high predictive power, we reasoned *VHL* function scores may be valuable for elucidating genetic mechanisms underlying unexplained genotype-phenotype associations. To illustrate this, we highlight two specific examples where SGE data nominate new mechanisms underlying genotype-phenotype relationships observed in humans.

Firstly, all nonsense SNVs located downstream of the C-terminal-most ɑ-helix (H4) scored neutrally by SGE (**Supplementary Fig. 9**). This suggests the last 12 AA of VHL are dispensable for HIF regulation. However, numerous indels have been identified in ccRCC samples between p.L201 and p.*214, calling the function of the region into question. Mapping indels observed in ccRCC with function scores shows a clear pattern, in which all ccRCC-associated indels downstream of the last LoF nonsense SNV share a common reading frame (**Fig. 5d**). These indels result in a 42-AA C-terminal extension (**Fig. 5e**), which is much longer than extensions created by indels in other reading frames and SNVs disrupting the normal termination codon. To validate this reading-frame specific effect, we analyzed CRISPR-induced indels in this region and confirmed growth defects are specific to indels leading to the 42-AA extension (**Fig. 5f**). This analysis indicates that between p.R200 and p.*214, frameshifting indels promote ccRCC development via a long C-terminal extension which impairs VHL’s ability to regulate HIF.

A second observation concerns c.264G>A (p.W88*). This variant introduces a stop codon downstream of p.M54 and has been shown to segregate with VHL disease in a family^24^. It is highly unusual among nonsense variants in that it was reported to cause type 2 disease marked by early-onset pheochromocytomas. Interestingly, c.264G>A did not score as LoF in our assay (function score = -0.16), in contrast to all other nonsense variants between p.M54 and p.L198, including c.263G>A, a nonsense variant at the same codon which causes type 1 disease (function score = -2.53; Supplementary Fig. 9). c.264G>A is also absent from cBioPortal ccRCCs. Therefore, the clinical evidence and our functional data are consistent in indicating c.264G>A preserves some ability to regulate HIF, despite being a nonsense variant.

One plausible mechanism is stop-codon readthrough. c.264G>A creates an opal codon, in context: 5’-<u>UGA</u>C. Opal codons followed by pyrimidines have been shown to be the most permissive to stop codon readthrough genome-wide^39^. In our SGE data, the next two highest scoring nonsense SNVs between p.M54 and p.L198 also created opal codons followed by pyrimidines (**Supplementary Fig. 9**). More functional evidence and clinical data will be needed to fully elucidate these mechanisms and to understand their importance in relation to cancer risk more broadly. Yet, these examples further illustrate how SGE data can resolve complex genotype-phenotype relationships by scoring variants along a spectrum of functional effects.

## DISCUSSION

Here, we applied a highly optimized SGE protocol to quantify the effects of nearly all SNVs across the coding sequence of *VHL*. Variant effects in the assay were dependent on HIF1A and predicted pathogenicity with high accuracy. Technological advances include i.) key optimizations to the SGE protocol leading to higher precision, ii) measurement of variant effects in isogenic lines to establish genetic dependencies, and iii) serial quantification of variants in mRNA revealing selective effects on expression. Combined with human phenotypic data, the SGE data constitute a mechanistically informative variant effect map of *VHL*.

Despite substantial improvements to the SGE protocol that reduce noise and promise to make the method applicable to more genes, limitations remain. Even with substantial optimization, we were unable to confidently score many variants introduced to the GC-rich, 5’-region of exon 1 where editing rates were lowest. However, the fact no variants tested prior to p.M54 scored as LoF suggests this region harbors few, if any, SNVs of clinical importance, a finding confirmed by the lack of pathogenic variants in this region reported in ClinVar and in ccRCC samples.

Towards the goal of identifying variants underlying ccRCC development, SGE performs remarkably well. The accuracy with which function scores distinguish pathogenic variants indicate the data may be used as strong evidence to support clinical decision-making.

Overall, only a small fraction of SNVs deemed ‘pathogenic’ or ‘likely pathogenic’ in ClinVar could not be distinguished from neutral variants using SGE data alone. The fact that none of these variants were observed in the ccRCC samples analyzed suggests some fraction may be misclassified. However, type 2 VHL disease variants have also been shown to mediate HIF-independent effects important for pheochromocytoma development^40, 41^. Such effects were not well represented in our HAP1-based assay, as we showed function scores were dependent on HIF1A expression. The fact that most pheo-predominant variants show a lesser degree of functional impairment (**Fig. 5c**) supports the conclusion that alleles leading to complete LoF do not typically manifest as type 2 VHL disease.

For patients harboring germline VUS in *VHL*, our data may provide valuable evidence in support of definitive classification^11^. With the recently proven efficacy of the HIF2ɑ-inhibitor belzutifan for treatment of VHL disease^14^, accurate classification of pathogenic variants will facilitate access to better care. Going beyond this, we anticipate our function scores may also help stratify patients with VHL disease by risk of specific tumors. Large cohorts of closely monitored patients will be needed to further explore how function scores correlate with clinical features of VHL disease, such as overall disease burden and likelihood of cancer progression. It will be intriguing to ask whether patients with germline variants that score lower via SGE are prone to develop ccRCC earlier, or to develop more distinct tumors.

Likewise, we envision our data being useful for adjudicating somatic *VHL* variants observed in tumor sequencing, especially in light of ongoing trials of belzutifan for sporadic ccRCC^42^. Across all cancers, LoF variants were predominantly seen in tumors previously associated with *VHL* mutations (**Fig. 4b-d**). Although *VHL* mutations occur much less frequently in renal cancers other than ccRCC, our data support their functional significance when present, a finding that suggests molecular profiling may identify additional RCCs that would respond favorably to treatments targeting HIF signaling. Conversely, more than one in ten *VHL* variants deemed “oncogenic” in cBioPortal scored neutrally by SGE, including several SNVs found in ccRCC. In these cases, treatments targeting HIF may provide little or no benefit.

Our analysis of both germline and somatic variants highlighted how orthologous lines of evidence can be leveraged to minimize errors in classification. This was evidenced by the perfect concordance between function scores and gold-standard ccRCC classifications supported by multiple data points (**Fig. 4f,g**). Owing to the gene’s compact size and the fact it is nearly always mutated in ccRCC, relatively few LoF variants were identified here that had not been previously observed in some form of clinical sequencing. For instance, among the *n* = 238 lowest scoring variants (function score less than -1.0), only 17 were altogether absent from ClinVar, cBioPortal, and VHLdb. Of these, all but one are created via transversion mutations, suggesting their rarity may be linked to lower mutation rates.

The RNA scores determined by SGE provide a means of examining the interplay between dosage effects at the mRNA level and functional output. This is an important relationship in the context of VHL disease because variants leading to complete loss of protein activity cause type 1 VHL disease. We show relatively few coding variants reduce mRNA dosage enough to impair VHL function in the assay. While we cannot preclude the clinical significance of variants impacting *VHL* mRNA levels to a lesser extent, particularly as they relate to recessive diseases such as congenital polycythemia^22, 43^, this finding suggests only non-coding variants with large effects on mRNA expression should warrant suspicion for a dominantly inherited cancer predisposition syndrome. Our RNA scores also provide additional value for identifying rare variants contributing to recessive phenotypes, such as c.222C>A. This synonymous SNV linked to recessive VHL deficiency^23^ had a marginally low function score of -0.40, but was highly depleted from the mRNA pool (RNA score = -4.5), consistent with its documented effect on splicing.

Collectively, this study and other recent implementations of SGE^29, 44, 45^ show how relatively simple assays reflective of cell-intrinsic effects can identify variants driving human disease with accuracies close to 100%. A key advantage of these approaches has been the ability to edit genes in their endogenous context. To date, though, SGE has only been performed in highly amenable cell systems. Therefore, a key challenge going forward will be to extend such approaches to more cell types and assays.

With continued genomic profiling of patients and tumors and increasing use of multiplex assays to systematically study thousands of variants at once^46^, our ability to map genotypes to phenotypes across a range of functional effects will continue to improve. In this context, we anticipate this analysis of *VHL* will prove highly valuable for clinical variant interpretation while also guiding future efforts to elucidate complex genotype-phenotype relationships across additional genes.

## FIGURES AND FIGURE LEGENDS

## Supplementary Figures

**Supplementary Fig. 1:**
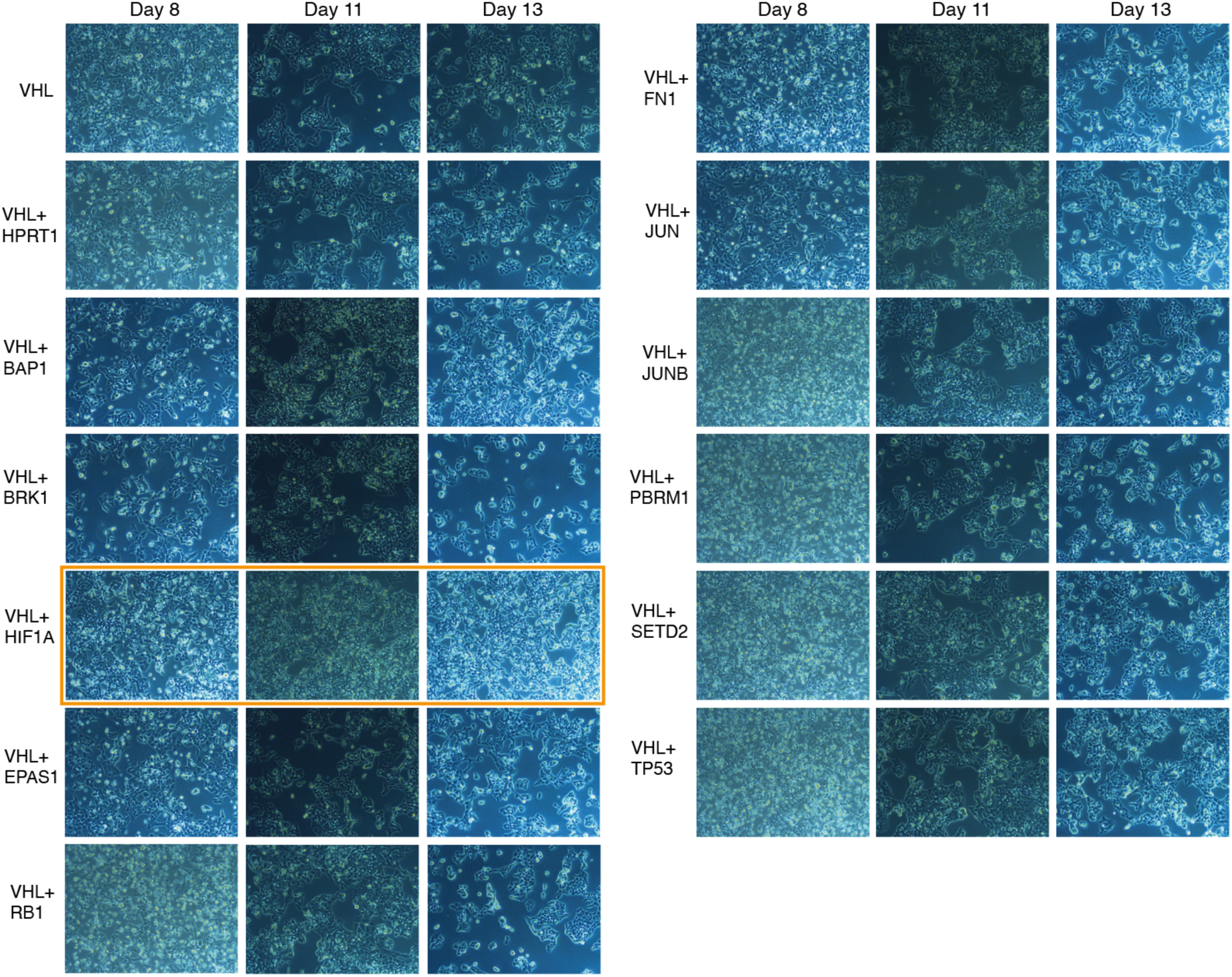
CRISPR-targeting of *HIF1A* rescues impaired HAP1 growth upon VHL depletion. HAP1 cells were transfected with plasmids expressing Cas9 and guide RNAs (gRNAs) targeting each of the genes indicated. Cell growth was assessed by light microscopy at three timepoints following transfection: day 8, day 11 and day 13. Double gRNA transfections were compared to targeting *VHL* alone or *VHL* plus a control gene (*HPRT1*). Co-targeting of *HIF1A* led to higher cell densities at later time points (orange box) compared to targeting *VHL* alone or *VHL* and other genes.

**Supplementary Fig. 2:**
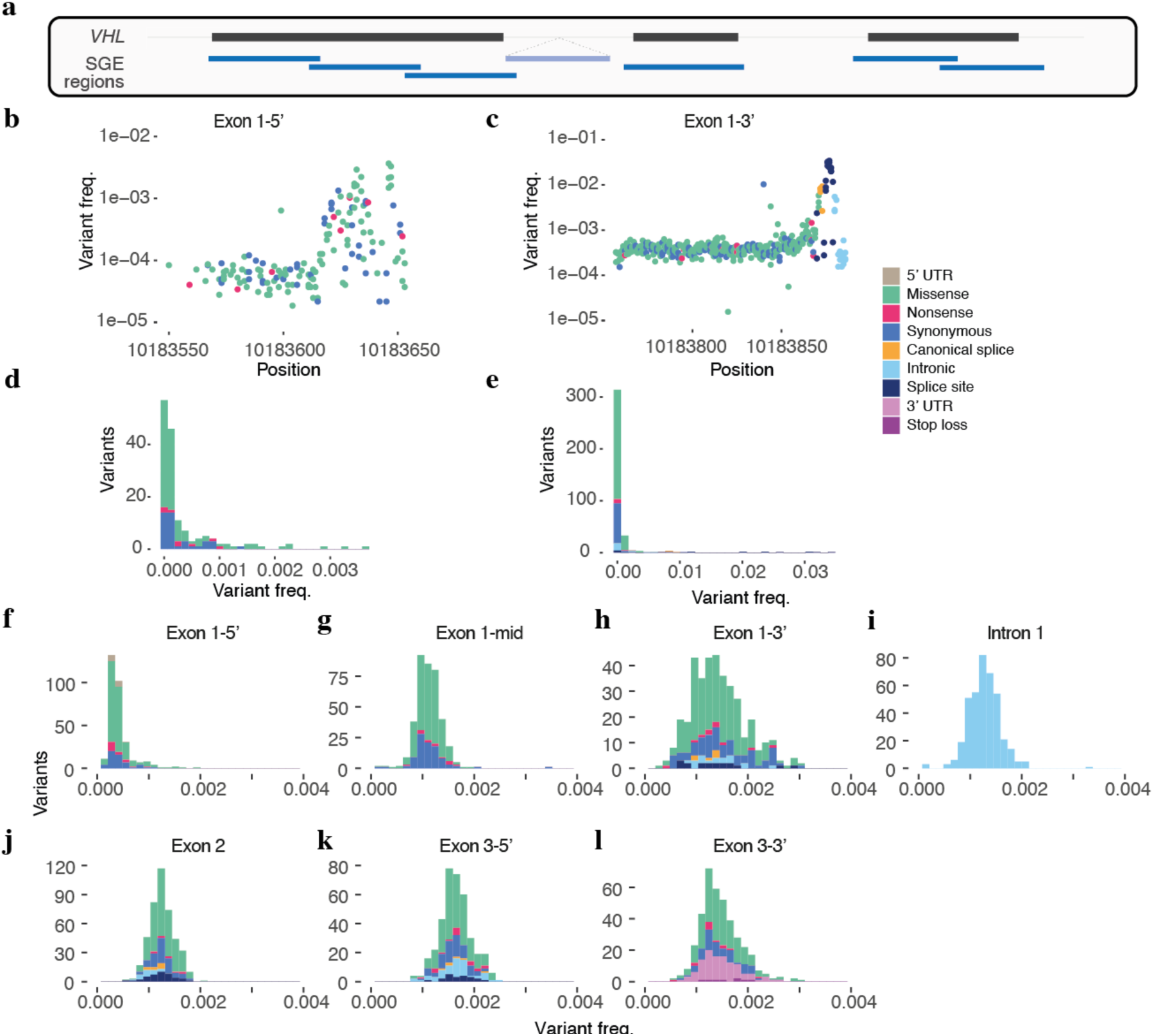
Optimizing SGE libraries to tile the complete *VHL* coding sequence. **a**, A schematic showing the seven SGE regions tiling across *VHL*. **b,c**, Frequency of SNVs plotted by position in the initial libraries for exon 1-5′ (**b**) and exon 1-3′ (**c**). **d**,**e**, Histograms of variant frequency for the initial libraries for exon 1-5′ (**d**) and exon 1-3′ (**e**). Based on these distributions, additional synonymous SNVs were added to final library designs. **f-l**, Frequency of SNVs in the final SGE libraries used for each region.

**Supplementary Fig. 3:**
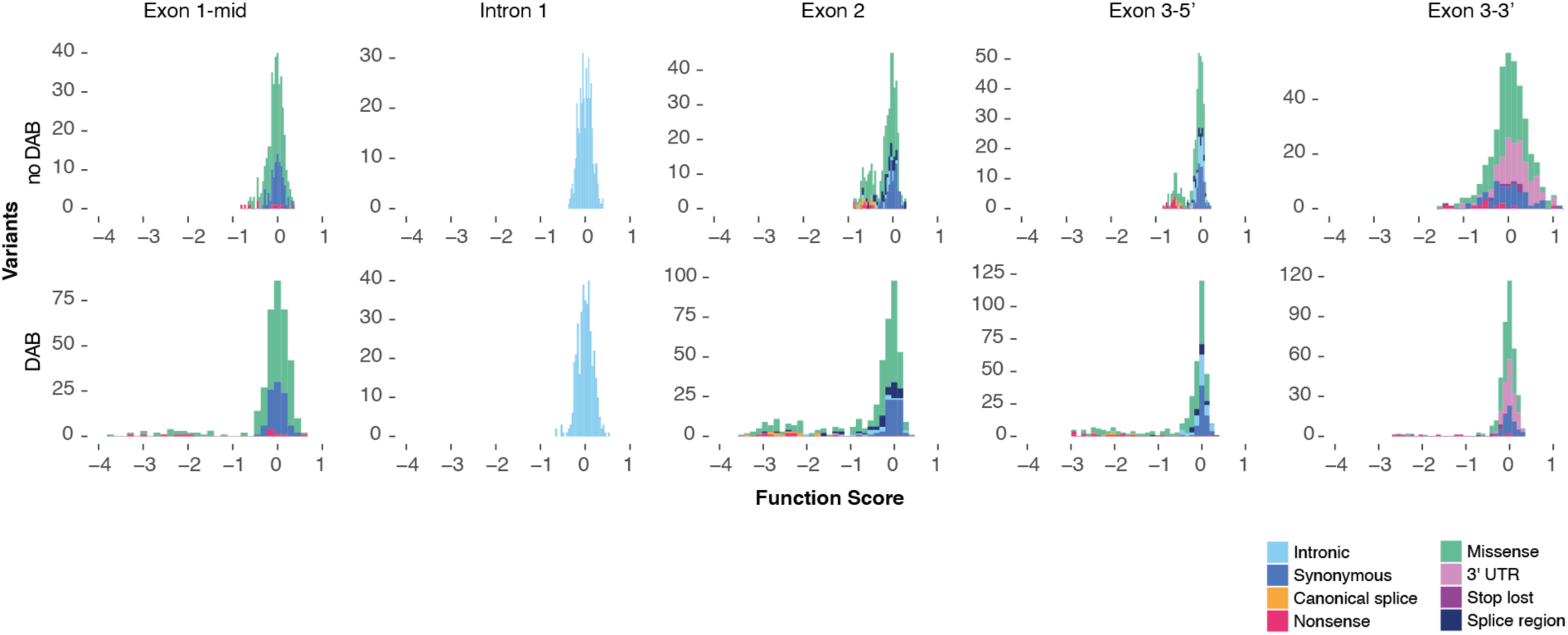
Addition of DAB to SGE experiments improves data quality. Histograms of function scores for regions where SGE was performed in normal HAP1 growth media (top) and media supplemented with 2.5 µM DAB (bottom). Function scores span a greater range when derived using DAB.

**Supplementary Fig. 4:**
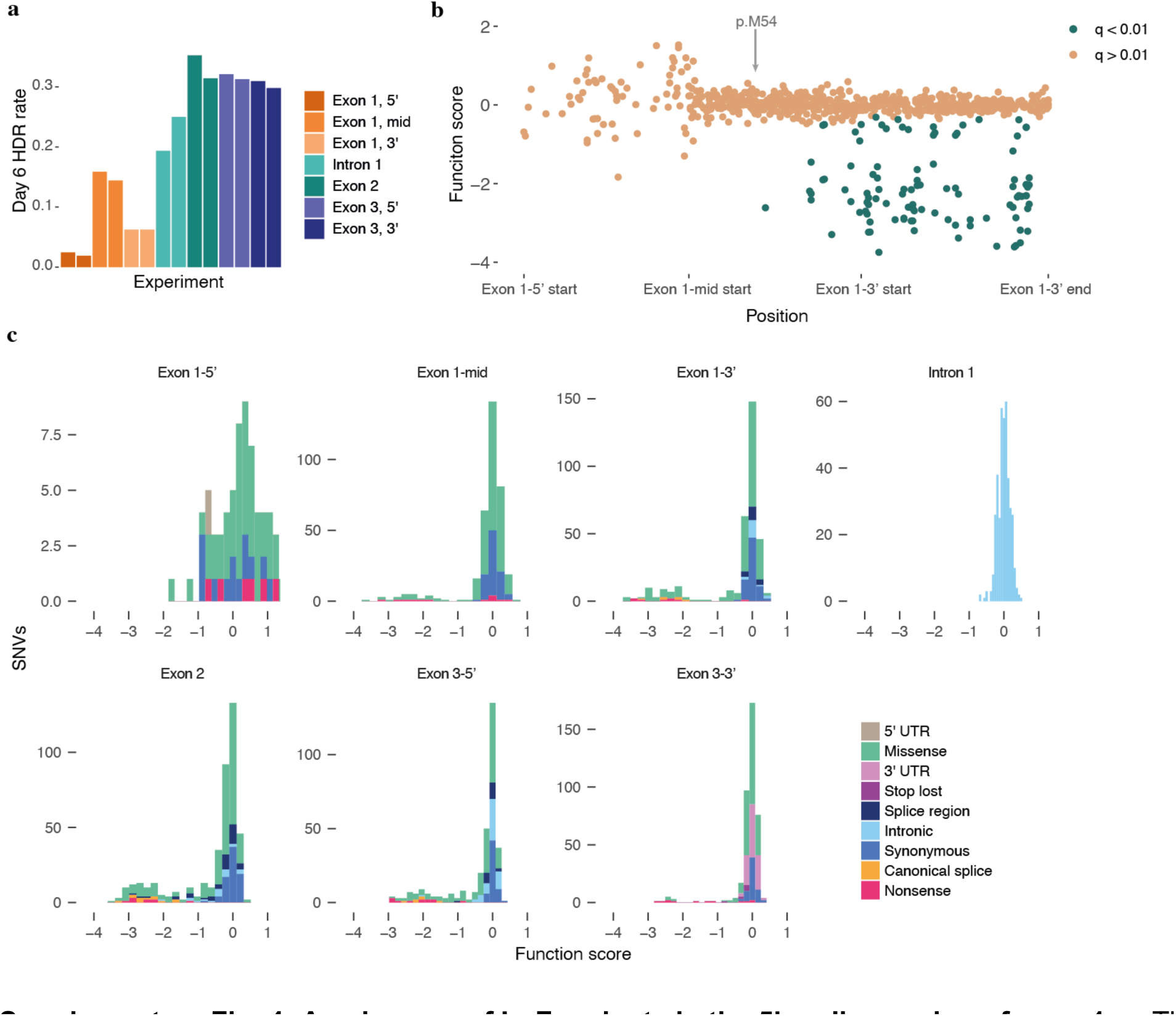
An absence of LoF variants in the 5’ coding region of exon 1. **a**, The rate of editing by HDR as measured by NGS is plotted for each replicate SGE experiment, sampled on day 6 post-transfection. **b,** Function scores for variants in exon 1 are plotted by genomic position and colored by q-value. Positions of the three different SGE regions tiling exon 1 are indicated on the x-axis. **c,** Histograms of function scores colored by mutation consequence are shown for each SGE region. Nonsense variants consistently score lowly across SGE regions, with the exception of the exon 1-5’ region.

**Supplementary Fig. 5:**
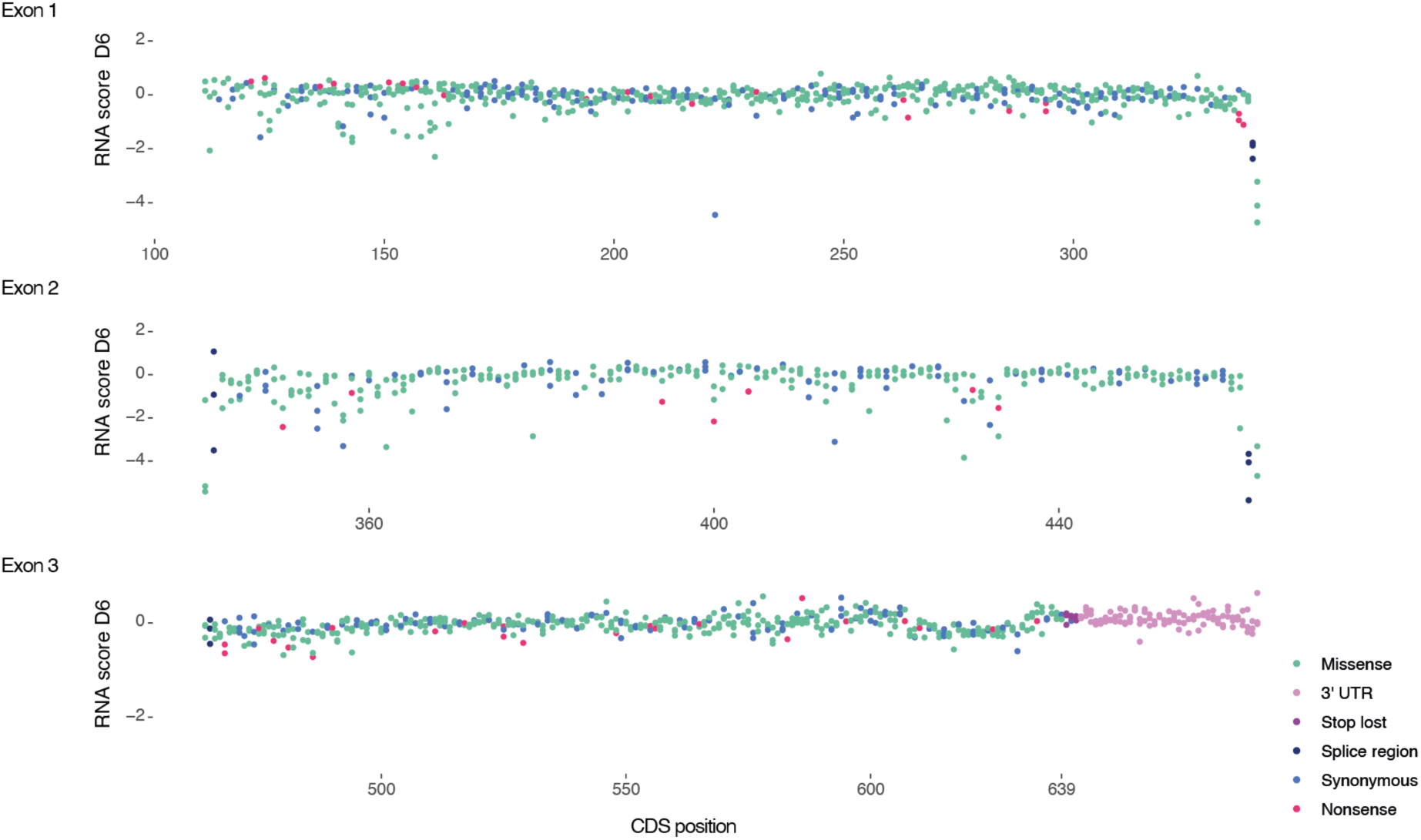
A map of RNA scores for *n* = 1,626 SNVs in *VHL*. RNA scores, defined as each SNV’s abundance in cDNA normalized to its abundance in gDNA, are plotted by transcript position. RNA scores shown are from samples harvested 6 days post-transfection.

**Supplementary Fig. 6:**
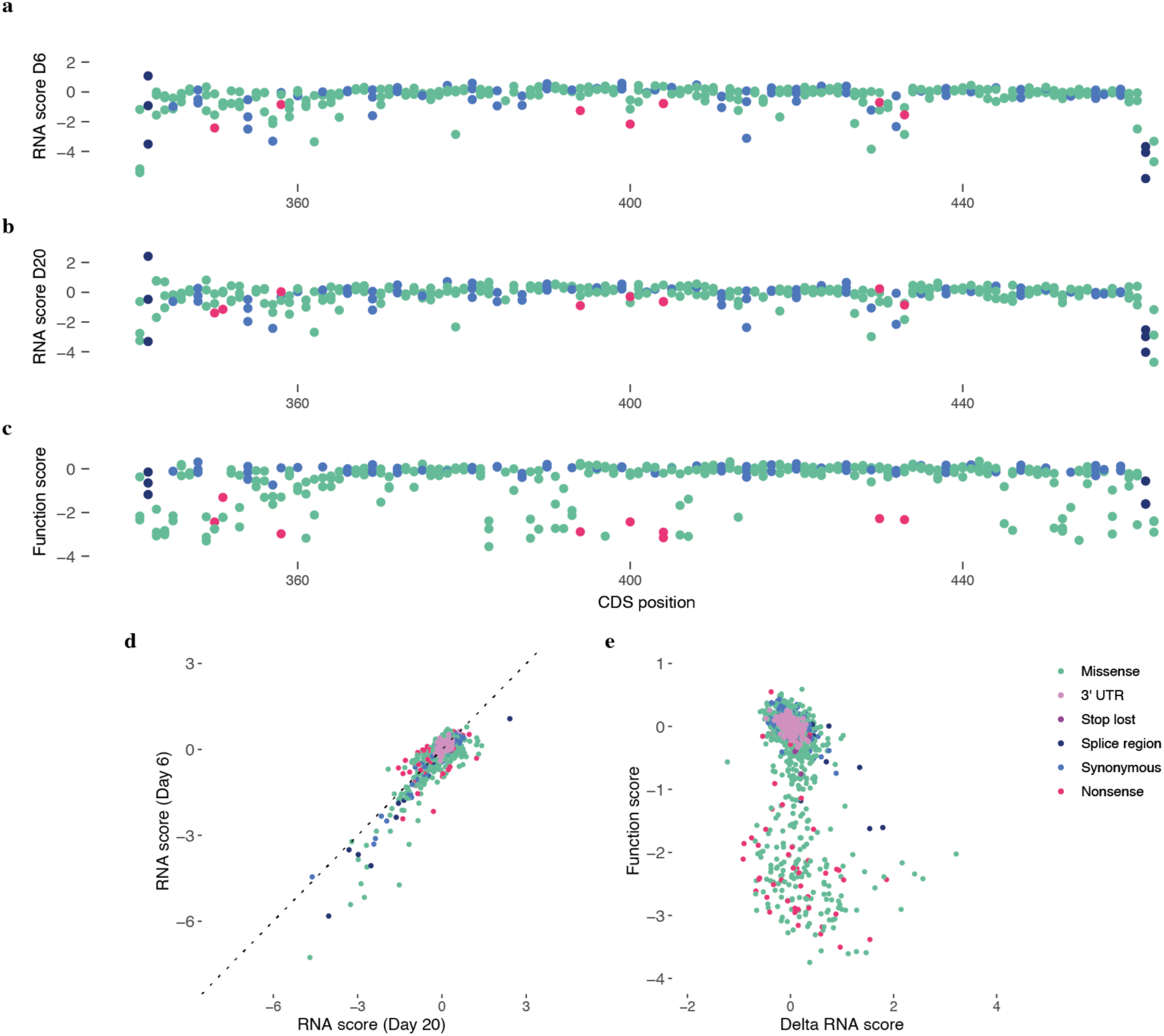
Comparing RNA scores across timepoints. **a-c**, Scores for *n* = 356 SNVs analyzed in exon 2 are plotted by transcript position. RNA scores are plotted for samples harvested 6 days post-transfection (**a**) and 20 days post-transfection (**b**). Function scores are plotted for the same set of exon 2 variants (**c**). **d**, RNA scores for all exonic variants (*n* = 1,626 SNVs) correlate broadly across timepoints (Pearson’s *R* = 0.86). Many variants with low RNA scores on day 6 have relatively higher RNA scores on day 20. (*y = x* plotted as a dashed line for reference.) **e**, The change in RNA score between day 6 and day 20 (delta RNA score) is plotted versus function score for the same set of SNVs. A subset of SNVs with low function scores have high delta RNA scores, indicating these variants were expressed in cells at higher levels on day 20 compared to day 6.

**Supplementary Fig. 7:**
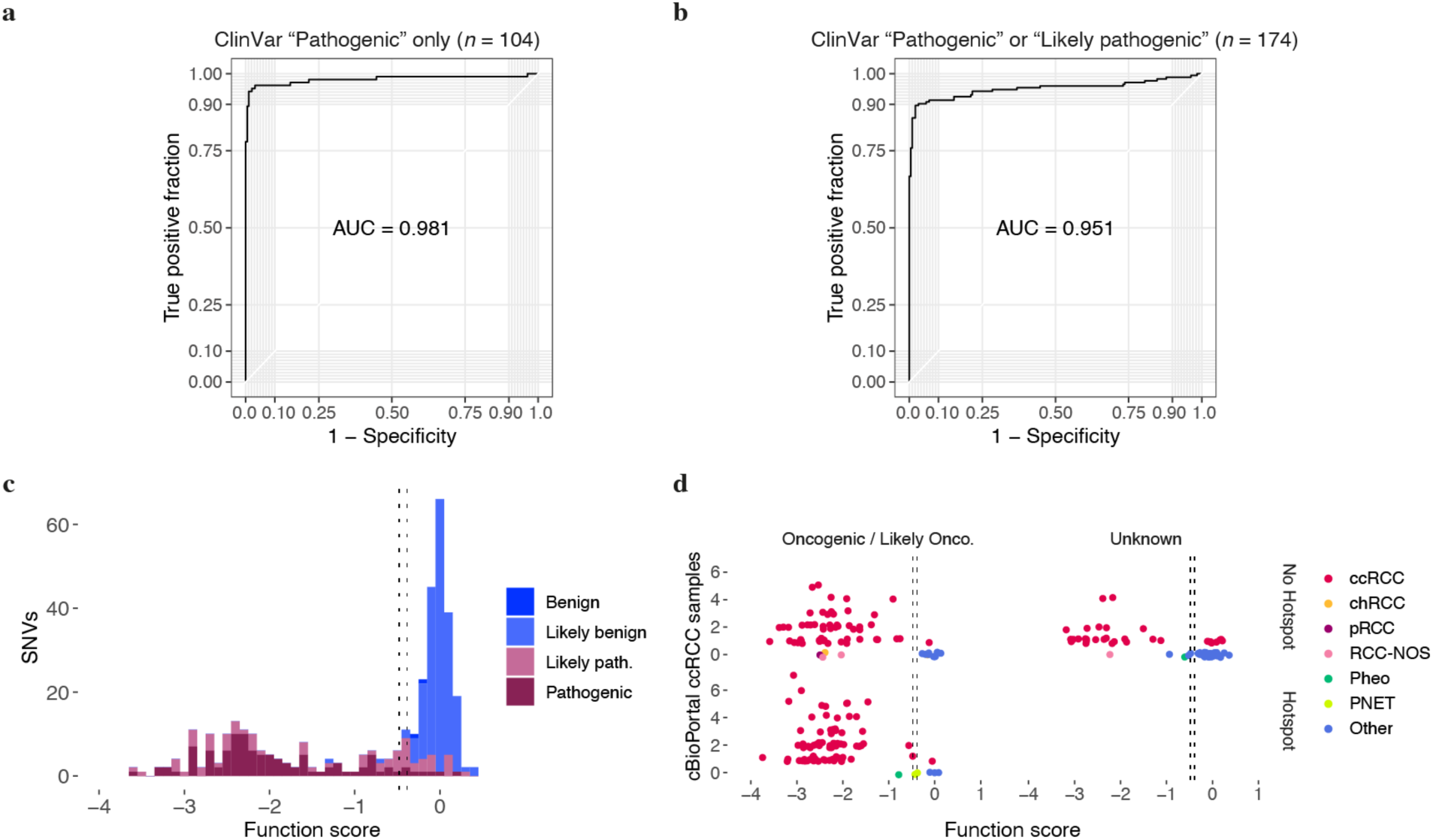
Function scores accurately predict pathogenicity of germline and somatic variants. **a,b**, Receiver operating characteristic (ROC) curves are shown for the classification of ClinVar variants using SGE function scores. “Pathogenic” SNVs (**a**) or “pathogenic” and “likely pathogenic” SNVs (**b**) were distinguished from *n* = 190 “Benign” or “Likely Benign” SNVs. **c**, The distribution of function scores is shown for ClinVar pathogenic and benign variants (*n* = 364 SNVs), with dashed lines indicating thresholds that distinguish the gold-standard ccRCC variants and neutral variants plotted in Fig. 4f. **d**, For each unique *VHL* SNV in cBioPortal (*n* = 233 SNVs), the function score is plotted versus the number of ccRCC samples in which the SNV was observed. Variants are split into separate plots by OncoKB annotation and mutational hotspot status, as reported in cBioPortal, with thresholds indicated as in (**c**).

**Supplementary Fig. 8:**
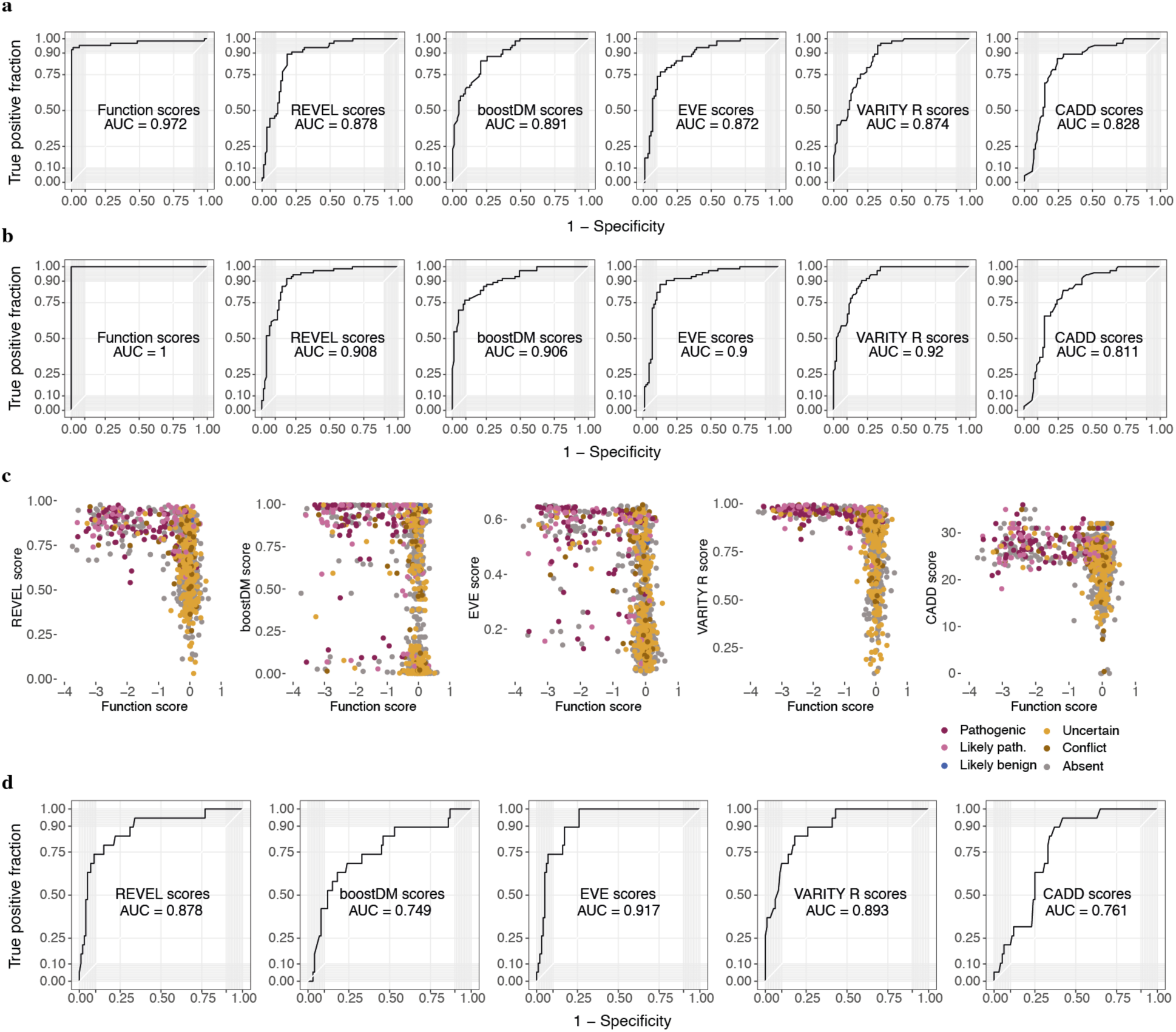
Function scores for missense variants outperform predictions from computational models. **a,b**, ROC curves indicate the performance of different metrics at distinguishing disease-associated missense variants in *VHL*. The metrics evaluated were SGE function scores, REVEL scores^5^, boostDM scores from the VHL-ccRCC model^8^, EVE scores^7^, VARITY R scores^9^, and CADD scores^25^. Missense SNVs were included if scored by all metrics (i.e. those present in SGE data from p.M54 to p.A207). In **a**, *n* = 65 missense variants deemed ‘pathogenic’ in ClinVar were distinguished from *n* = 87 missense SNVs deemed neutral (as in Fig. 4g). In **b,** missense variants present in the gold-standard set of ccRCC-associated SNVs (*n* = 73) were classified against the same neutral set of variants as in (**a**). **c**, Function scores for *n* = 953 missense SNVs are plotted versus scores from each computational predictor, colored by ClinVar status. **d**, Function scores were used to define two sets of unseen variants (i.e. those absent from ClinVar, cBioPortal, population sequencing, and VHLdb). Each metric was assessed on its ability to distinguish unseen missense SNVs with function scores below -0.479 (*n* = 19) from the set of missense SNVs with function scores closest to 0 (*n* = 100).

**Supplementary Fig. 9:**
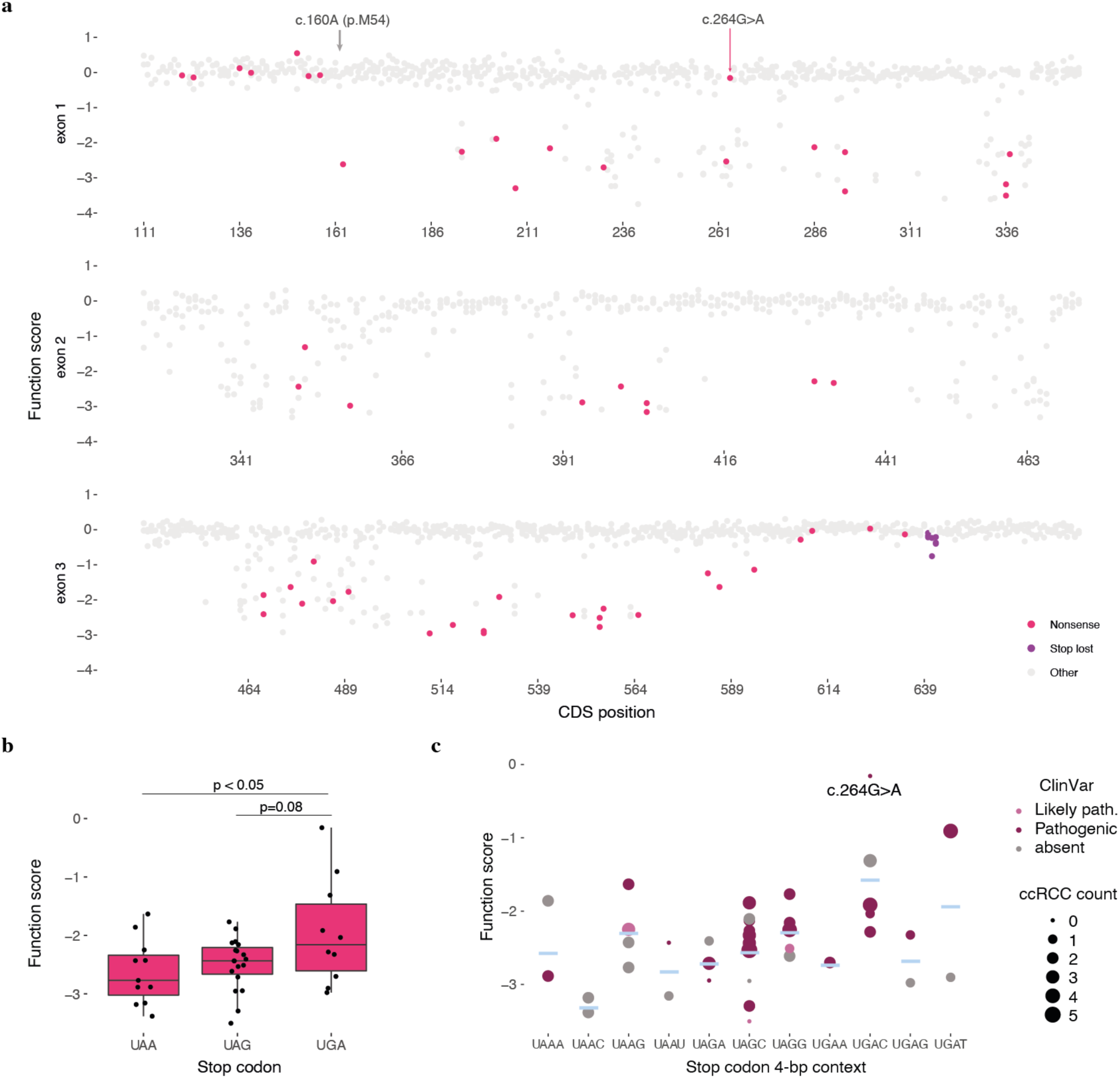
Functional effects of nonsense variants in relation to position and stop codon context. **a**, Function scores are plotted by position in *VHL* and colored to highlight nonsense variants. All nonsense SNVs tested between c.160A and c.601C scored as LoF, except for c.264G>A, a variant associated with type 2 VHL disease^24^. **b,c**, Function scores for nonsense variants between c.160 and c.601 are plotted by termination codon (**b**) and 4-bp termination codon context (**c**). Differences between function scores by termination codon were tested using a one-way ANOVA. (Boxplot: center line, median; box limits, upper and lower quartiles; whiskers, 1.5x interquartile range; all points shown.) In **c**, the blue line indicates the mean score for each stop codon context and the size of each dot corresponds to the number of times the SNV has been reported in cBioPortal ccRCC samples.

## METHODS

### HDR library design and cloning

SGE experiments were designed as described previously^29^ using the full-length *VHL* transcript (ENTS000000256474, CCDS2597). Six SGE regions of length 128-146 bp were designed to cover the entire coding sequence of *VHL*. Regions for mutagenesis included all exonic sequences, as well as exon-proximal regions of introns, extending up to 27 bp. An additional SGE region was designed in intron 1 to cover positions previously associated with splicing alterations^43^.

Oligonucleotide libraries were designed for each SGE region. First, synonymous substitutions were designed at each of two CRISPR protospacer adjacent motif (PAM) sequences in the region. The purpose of these synonymous PAM edits was to prevent Cas9 from re-cutting the genome following the HDR and to distinguish true HDR-derived SNVs in NGS analysis. These substitutions were included in the template sequence to which all possible SNVs were introduced to make the SGE library (**Supplementary Table 2**).

Sequences were generated for every possible single nucleotide substitution within each SGE region. Unedited genomic sequence was appended to ends of each oligo to facilitate region-specific PCR amplification and cloning into homology vectors. The final designed oligonucleotide libraries contained molecules representing all possible SNVs within each SGE region.

For two of the SGE regions (exon 1-5′ and exon 1-3′), additional synonymous substitutions were added in a second library design round to reduce repeats and high-GC sequence content, thereby improving uniformity. All oligonucleotide libraries were synthesized (Twist Bioscience) as a single oligonucleotide pool and resuspended at 5 ng/μL. Primers complementary to the adapter sequences in the oligonucleotide libraries were designed to amplify variant libraries from the oligonucleotide pool for each SGE region. KAPA HiFi ReadyMix (Roche) was used to perform PCRs to amplify libraries in 25 μL reactions, using 500 pg of oligonucleotide pool as DNA template for each reaction. All PCR reactions were monitored with real-time (RT-)PCR by spiking SYBR green (Thermo Fisher) into the reactions at 0.4x final concentration. Cycling was stopped upon completion of amplification.

For each SGE region, a vector containing homology arms was generated by PCR-amplification of HAP1 gDNA and cloning of products into a linearized pUC19 backbone (InFusion, Takara Bioscience). Primers for these PCRs were designed such that homology arms would be between 200 bp and 1300 bp. The homology arm plasmids were subsequently linearized by inverse PCR using primers with 15-20 bp of overlap with the corresponding adapter primers used to amplify the oligo libraries. In the linearization PCRs, 10 pg of each plasmid containing unedited gDNA was used as template in each 50 μL reaction, and the resulting products were subject to DpnI digestion for 1 hour at 37 ℃.

Amplified oligo pools and PCR-linearized homology arm vectors were purified using AMPure XP beads (Beckman Coulter). To generate final HDR libraries, the amplified oligo libraries were cloned into the linearized pUC19-homology arm vectors. In each cloning reaction, approximately 50 ng of insert and 65 ng of vector were used. The resulting products were transformed into Stellar Competent *E. coli* (Takara Bioscience). 1% of the transformed cells were plated on ampicillin-agar plates to ensure adequate transformation efficiency (at least 10-fold library coverage), and the remaining cells were cultured overnight at 37 ℃ in 150 mL of LB with carbenicillin (100 ug/ml).

### CRISPR gRNA design and cloning

Target sites for *S. pyogenes* Cas9 were designed such that DNA cleavage would occur within coding sequences of *VHL* and such that synonymous substitutions could be used to disrupt the target PAM sequence, where possible. When multiple gRNAs were suitable for a given SGE region, gRNAs were chosen to have minimal predicted off-target activity (Benchling).

To anneal and clone the gRNAs, a previously described procedure was followed, including treatment of the ligation reaction with PlasmidSafe DNase (Lucigen)^49^. Briefly, annealing and phosphorylation of gRNA-specific oligonucleotides was carried out in a thermocycler as follows: phosphorylate oligonucleotides with polynucleotide kinase at 37 ℃ for 30 minutes, denature at 95

℃ for 5 minutes, slowly cool to 25 ℃ at 5 ℃ per minute. Digested plasmid to co-express a gRNA and Cas9 with a puromycin resistance cassette (pX459; Addgene) was purified from a 1% agarose gel using a Gel and PCR Clean-up Kit (Macherey-Nagel) according to the manufacturer’s protocol. The ligated reactions were transformed into Stellar Competent *E. coli* cells (Takara Bioscience). Sanger sequencing (Genewiz) was used to confirm correct gRNA sequences. Sequence-verified plasmids were purified with the ZymoPure Maxiprep kit (ZymoResearch) and eluted in nuclease-free water (Invitrogen).

### Tissue culture: Subculture routine

To perform SGE, HAP1 cells with a *LIG4* frameshifting deletion (HAP1-LIG4-KO) were used as described^29^. Cells were cultured with Iscove’s Modified Dulbecco’s Medium (IMDM) containing L-glutamine and 25 nM HEPES (Gibco) supplemented with 10% FBS (Gibco) and 1% Pen/Strep (Gibco). Cells were thawed one week prior to transfection to allow for recovery and expansion and maintained at under 80% confluency. Each passage, cells were washed twice with 1X DPBS (Gibco), trypsinized with 0.25% trypsin-EDTA (Gibco), resuspended in media, centrifuged at 300 x *g* for 5 minutes, and resuspended. No fewer than 6 million cells were re-plated, splitting either 1:5 across 2 days or 1:10 across 3 days. Apart from initial SGE experiments performed without DAB, DAB (Stratech) was freshly added to the media each passage at 2.5 uM.

### Tissue culture: SGE experiments

One day prior to transfection, 15 million cells were seeded on a 10-cm dish in 10 mL of media. For each SGE region, cells were co-transfected with 30 μg of pX459 plasmid expressing a region-specific gRNA and 10 μg of the corresponding HDR library. Xfect (Takara Bioscience) transfection reagent was used, and the manufacturer’s protocol was followed with the following modifications:

1. On the day of transfection (day 0), cells were 80-90% confluent.
2. 0.6 μL of Xfect polymer was used for each μg of DNA.
3. 40 μg of total plasmid DNA was added per transfection of each 10-cm dish.
4. The final volume of transfection buffer, DNA, and Xfect polymer added to cells was 800 μL.

Two replicate transfections were performed for each SGE region, with plates incubated overnight. On day 1, cells were washed and transferred to a 15-cm dish and resuspended in fresh media. Puromycin (Cayman Chemical) was added to a final concentration of 1 μg/mL to select for transfected cells. During puromycin selection, DAB was not added to media. On day 4, the cells were passaged to fresh media supplemented with DAB, without puromycin. Cells were harvested on days 6, 13 and 20. At each time point, 10 million cells were collected as cell pellets and stored at -80 ℃.

Negative control samples for each SGE region were co-transfected with the HDR library and a pX459 plasmid containing a gRNA targeting *HPRT1*. The same transfection conditions were used but scaled proportionally to a single well of a 6-well plate. Negative control samples were harvested on day 6.

### Tissue culture: Generation of monoclonal cell lines

HAP1-LIG4-KO cells were transfected with a pX459 plasmid targeting *HIF1A* as described above. Cells were counted with the Vi-Cell XR Analyzer (Beckman) and diluted to 0.8 cells per 100 μL. 100 μL were aliquoted into each well of a 96-well plate. After one week, light microscopy was used to identify wells with a single growing clone. 10 single-cell clones were expanded and maintained. gDNA was extracted from each line (DNeasy Blood & Tissue Mini kit, Qiagen), PCR-amplified, and Sanger sequenced. A HIF1A-knockout clone containing a 7-bp deletion and a single base substitution was selected for use as the HIF1A-knockout line.

### Tissue culture: Essentiality testing

To test the essentiality of *VHL* in HAP1, a pX459 vector expressing a gRNA to target exon 2 of *VHL* was transfected alone (day 0). Transfections were performed as described above, transfecting cells with 5 μg of DNA in a single well of a 6-well plate, in replicate. Samples were harvested on day 6 and day 13 to analyze selection on editing outcomes.

### gDNA and RNA extraction

Harvested cells were stored as pellets at -80 ℃ until ready to proceed. QIAshredder columns (Qiagen) were used to homogenize cells and the AllPrep DNA/RNA kit (Qiagen) was used to purify RNA and gDNA. Procedures were followed as per the manufacturer’s protocol. Concentrations were measured using Nanodrop UV spectrometry for RNA, and the Qubit BR Kit for gDNA (Thermo Fisher).

### RNA preparation for sequencing

cDNA was generated from 5 μg of RNA per sample using the SuperScript IV First-Strand Synthesis System (Invitrogen). A *VHL*-specific primer complementary to the gene’s 3’ UTR was used for priming the reaction, and the manufacturer’s protocol followed. cDNA samples were subsequently prepared for sequencing starting with PCR 2 below.

### PCR 1: amplifying gDNA

All primers used are provided in **Supplementary Table 2**. For each reaction, 2 μg of gDNA was PCR-amplified using at least one primer to target gDNA outside of the homology arms included in HDR libraries, therefore making the reaction specific to gDNA. Annealing temperatures and cycling times were optimized for each SGE region in advance using gDNA from unedited HAP1 cells. Up to eight 100 μL reactions were carried out for each sample, or fewer depending on gDNA yield. Magnesium chloride was added to a final concentration of 5 mM and KAPA HiFi 2x ReadyMix (Roche) was used. The reactions for each sample were pooled together and correct products were confirmed using gel electrophoresis. PCR products were purified using AMPure XP (Beckman Coulter) according to protocol. The same procedure was carried out for negative control transfections and SGE gDNA samples from each timepoint (days 6, 13, and 20).

### PCR 2: adding Nextera adapters

A second, nested PCR for each sample was carried out using 1 μL of the purified product from PCR 1. For this reaction, primers were designed for each SGE region to produce amplicons compatible with Illumina-based NGS sequencing, such that 300-cycle kits would provide end-to-end coverage of amplicons. Primers added Nextera sequencing adapters to the amplicons to enable paired-end sequencing. Products were verified using gel electrophoresis and purified using AMPure XP (Beckman Coulter). The same reaction was also performed to prepare each HDR library for sequencing. cDNA samples were similarly amplified using cDNA-specific primers designed to yield amplicons spanning SGE regions in coding sequence, as well as at least 1 exon junction.

### Indexing and sequencing

Each sample was dual-indexed by PCR using custom indexes and purified using AMPure XP. Samples were quantified using the Qubit HS kit (Invitrogen). Illumina guidelines were followed to dilute and denature samples before sequencing using an Illumina NextSeq (mid-or high-output 300 cycle kits). Approximately 5 million reads were allocated for each SGE gDNA sample, except for samples from high-GC regions (exon 1-5′ and exon 1-3′), which were allocated 10 million reads. 1 million, 2 million, and 3 million reads were allocated for the negative control, library, and RNA samples, respectively. 20-30% PhiX control (Illumina) was included in each sequencing run.

### Processing SGE sequencing data

A custom pipeline was used to process all sequencing data to variant counts. Bcl2fastq2 (Illumina) was used to create demultiplexed fastq files. Paired-end reads were then adapter-trimmed and merged if the overlapping sequence between reads matched perfectly (SeqPrep). Merged reads containing N bases were removed, prior to performing global alignment with needle-all (EMBOSS package) against a reference amplicon for each SGE region. Resulting sam files were processed using a series of custom scripts to analyze indel events from CIGAR strings, to count SNV frequencies across samples, and to annotate variants by genomic position with data from CADD v1.6 (hg19)^25^. For calculating function scores, reads for each variant were only included if at least 1 silent PAM edit in the HDR library design was present in the read and no other variants were seen. This filter is designed to remove reads resulting from 1-bp substitutions to unedited templates arising via error during PCR or sequencing.

### Deriving function scores for SNVs

In a series of quality filters, SNVs were removed from analysis if:

1. The variant’s frequency in the HDR library was less than 1.0×10^-4^.
2. The variant’s day 6 frequency in either replicate was less than 1.0×10^-5^.
3. The day 20 to day 6 ratio was highly discordant across replicates (i.e. a difference in log_2_ ratio greater than 1.5, unless both ratios were less than -1.0.
4. The day 13 to day 6 ratio was markedly different from the day 20 to day 13 ratio (i.e. a difference in log_2_ ratio greater than 2.0, unless both ratios were less than -0.5).
5. Estimates of sequencing error from negative control samples suggested the ratio of SNV observations arising from sequencing error to be greater than 0.5.
6. In instances where a variant was engineered in the same codon as a synonymous PAM edit, the variant was excluded if the resulting amino acid change was different from what it would be without the PAM edit also present.

*To calculate function scores, the mean log_2_ day 20 to day 6 ratio for each variant across replicates was normalized to the median synonymous variant in the sample. Final function scores were then normalized across exons using the range of effect sizes observed for synonymous and nonsense variants. Explicitly, scores for each SGE region were scaled linearly such that the median nonsense variant for the region equaled the global median nonsense variant and the median synonymous variant scaled to 0. Only nonsense variants between p.54 and p.198 were used to perform this scaling (i.e. those expected to result in LoF).

### Calculating RNA scores

RNA scores were determined for exonic variants by using variant frequencies in targeted sequencing of *VHL* cDNA. Similar to variant frequencies derived from gDNA, variant frequencies were calculated only from reads in which both the variant and at least one PAM edit indicative of HDR was present. RNA reads were only included in scoring if the sequence on either side of the target exon matched the reference transcript perfectly (i.e. reads from canonically spliced transcripts). RNA scores for day 6 and day 20 were derived by normalizing each coding SNV’s frequency in RNA to its frequency in the gDNA sampled from the same time point. Non-coding variants were not assigned RNA scores because such variants are not detectable in amplicon sequencing of mRNA. RNA scores were also not generated for variants in the exon 1-5’ SGE region.

### Producing a single set of scores

For variants covered in multiple experiments (e.g. by overlapping SGE regions), final function scores and RNA scores were determined as follows:

1. For variants scored in both SGE regions exon 1-5’ and exon 1-mid, the scores from the exon 1-mid experiment were used.
2. For variants scored in both SGE regions exon 1-mid and exon 1-3’, the mean of the scores was used as the final score.
3. For variants scored in both SGE regions exon 3-5’ and exon 3-3’, the mean of the scores was used as the final score.

### Analysis of indels

To analyze selection on indel events, CIGAR strings from each timepoint sampled were generated via global alignment. Similar to function scores for SNVs, “indel scores” for each unique editing outcome (i.e. CIGAR string) were derived by dividing its day 13 frequency by its day 6 frequency. Scores were log2-normalized and averaged across replicates. Only editing outcomes with no more than a single indel and at least 100 bp of matching sequence on the 5’ end of the alignment were included. Analysis was restricted to outcomes observed in more than 0.1% of reads.

### Statistical testing

Statistical tests were performed in R. Unless otherwise indicated, all tests were two-sided.

To determine whether a given function score was significant, a null model was fit for each SGE region using all synonymous variants. In regions where RNA scores were included, synonymous variants were excluded from the null group if they had RNA scores (day 6) less than -1.0 (corresponding to a 50% reduction in mRNA levels). For experiments in which RNA scores were not derived (those without DAB, those performed in HIF1A-KO cells, and those performed in the exon 1-5’ SGE region), all synonymous SNVs were included in null models. For the intron 1 SGE region, all SNVs were included in the null model due to the lack of synonymous SNVs. P-values were calculated using the ‘pnorm’ function in R, and adjusted using ‘p.adjust’ (Benjamini-Hochberg method) to produce q-values. A false discovery rate of 0.01 was used as a cut-off for defining LoF SNVs.

### DepMap analysis

DepMap CRISPR screening data for *VHL* was downloaded via https://depmap.org/portal/. The “Primary.Disease” classification was used to define lines of kidney origin, and lines with a *VHL* mutation defined as either “Hotspot”, “Damaging”, or “Other non-conserving” were labeled mutant for *VHL*.

### Mapping variant effects to the VHL structure

The crystal structure of VHL in complex with ELOC, ELOB, and the HIF1A target peptide (PDB: 1LM8)^47^ was imported to PyMol. Average function scores of missense SNVs at each residue from p.M54 were calculated, excluding variants with RNA scores below -2.0. The VHL structure was colored by average function score to identify positions intolerant to missense mutations.

Alphafold’s colab notebook was used to model a structure for VHL with the 42-AA extension at the C-terminus: https://colab.research.google.com/github/deepmind/alphafold/blob/main/notebooks/AlphaFold.ip ynb.

### ClinVar and cBioPortal analyses

*VHL* entries with at least a 1-star assertion criteria rating in ClinVar were obtained on 04 May 2023. ClinVar annotations were simplified such that variants classified as both “pathogenic” and “likely pathogenic” by separate contributors were labeled “likely pathogenic” and variants deemed both “benign” and “likely benign” were labeled “likely benign”.

cBioPortal data was accessed on 02 October 2022 via https://www.cbioportal.org/. Mutation data from the “curated set of non-redundant studies” was exported by querying for samples with *VHL* mutations by gene symbol. Data were parsed using a custom script to sum the number of times each variant was seen across each cancer type. For analyses involving allele frequencies in tumors, each independent sample was plotted (i.e. a single variant present in multiple independent samples was represented multiple times). ccRCC samples “with sarcomatoid features” were grouped with all ccRCC samples. To assign a single cancer type to color variants seen in multiple cancers, preference was given to cancer types associated with *VHL* mutations, by order of priority: ccRCC, pheochromocytoma, PNET, chromophobe RCC, papillary RCC, RCC not otherwise specified, and other). Using this logic, variants were only colored “other” if they were not seen in tumors associated with *VHL* mutations.

### Population sequencing analysis

UK Biobank variants in *VHL* were ascertained via GeneBass (https://app.genebass.org/), which includes data from whole exome sequencing of 394,841 individuals (7 June 2022 data release). TOPMed (freeze 8) *VHL* variants were obtained (https://bravo.sph.umich.edu/freeze8/hg38/), as were non-overlapping gnomAD v2 and v3 data sets (https://gnomad.broadinstitute.org/). A combined population allele count was defined as the sum of gnomAD v2 and v3 allele counts, TOPMed heterozygous allele counts, and UK Biobank allele counts.

### VHLdb analysis

Mutation data from VHLdb^21^ was downloaded on 28 July 2022 and parsed using a custom python script to count features of entries for each variant. Where there was only 1 possible match to the SGE data set, the nucleotide change of an entry was inferred from the amino acid substitution if only the latter was provided. When the provided nucleotide and protein changes were discordant, the entry was removed, as were entries flagged “needs revision”. The disease descriptor “Von Hippel-Lindau syndrome: Pheochromocytoma phenotype not described” was replaced with “Von Hippel-Lindau syndrome: phenotype not described”, and the descriptor “absence of pheochromocytoma” was ignored in order to quantify pheochromocytoma reports.

“Pathogenic” and “likely pathogenic” variants in ClinVar, excluding variants with recessive phenotypes and those seen in two or more population control individuals, were assigned to categories of “type 1”, “pheo-predominant”, or “type unclear” by searching VHLdb for indicators of disease type and tumor type. Variants only associated with type 1 disease or ccRCC (without pheochromocytoma) were deemed “type 1”. Variants associated only with type 2 disease or with more pheochromocytoma entries than ccRCC entries were deemed “pheo-predominant”. The remaining “pathogenic” and “likely pathogenic” variants either had mixed associations, lacked phenotypic information, or were explicitly type 2B variants associated with high risk for both ccRCC and pheochromocytoma. All were deemed “type unclear”.

### Comparisons to computational predictions

CADD scores (v1.6)^25^ and annotations were obtained for all SNVs assayed (https://cadd.gs.washington.edu/download). The following scores were obtained for missense variants: REVEL scores^5^ were obtained on 01 September 2022 (https://sites.google.com/site/revelgenomics/downloads); scores from the boostDM model for *VHL* variants in “Renal Clear Cell Carcinoma” were obtained on 12 Aug 2022 (https://www.intogen.org/boostdm/search?ttype=RCCC&gene=VHL); EVE scores were obtained on 15 Aug 2022 (https://evemodel.org/); VARITY scores were obtained on 16 Aug 2022 (http://varity.varianteffect.org/).

For comparisons of missense variant effects, only variants for which all metrics provided a score were used. This excluded variants prior to p.M54 and variants after p.A207 due to an absence of EVE scores. ROC curves were generated in R using the ‘geom_roc’ function.

## DATA AVAILABILITY

All function scores and RNA scores derived are included in Supplementary Table 1, as well as raw NGS read counts per variant. Function scores will be made publicly available and deposited to MAVE-DB^50^ at time of publication. Fastq files will be made publicly available at time of publication (European Nucleotide Archive accession: PRJEB62650).

## CODE AVAILABILITY

All code used in this study is available on GitHub at https://github.com/TheGenomeLab/VHL-SGE.

## ACKNOWLEDGEMENTS AND FUNDING

We thank Robert Goldstone and Mike Gavrieldes for assisting with data access, Jerome Nicod, Marg Crawford, and the Advanced Sequencing STP for performing sequencing, Erik Sahai and David Balchin for helpful insights and feedback on the manuscript, Andrew Waters and David Adams for experimental advice, Peter Radcliffe and lab for discussion of VHL-related research, and the Cell Services STP for maintenance of cell lines.

This work was supported by the Francis Crick Institute which receives its core funding from Cancer Research UK (CC2190), the UK Medical Research Council (CC2190), and the Wellcome Trust (CC2190), and by funding from the VHL Foundation UK/Ireland (PRJ20071). For the purpose of Open Access, the authors have applied a CC BY public copyright license to any Author Accepted Manuscript version arising from this submission.

## SUPPLEMENTARY INFORMATION

**Supplementary Table 1. SGE function scores and RNA scores for 2,268 *VHL* SNVs.**

**Supplementary Table 2. Oligonucleotides used to perform SGE of *VHL*.**

